# A unified framework for causal gene regulatory network inference grounded in orthogonal molecular evidence

**DOI:** 10.64898/2026.04.28.721354

**Authors:** Ruihao Li, William K. M. Lai, Franklin B. Pugh

## Abstract

Gene regulatory networks (GRNs) govern gene expression, cellular differentiation, and stable transcriptional states. Yet inferring GRNs that integrate molecular regulatory mechanisms and reproduce transcriptional states as stable outcomes remains a central challenge. Here we present SETIA, a framework that infers GRNs whose explicit dynamical models reproduce transcriptional profiles as one or more stable states across conditions. Applied to RNA–seq data from wild–type and transcription factor knockout strains in *Saccharomyces cerevisiae*, SETIA infers GRNs that accurately reproduce held–out transcriptional states in cross–validation experiments. Incorporating TF–promoter binding and protein–protein interaction priors, SETIA yields GRNs ranging from mechanistically grounded architectures to flexible models that capture indirect regulatory influences. SETIA reveals that gene expression organizes into discrete stable states that represent distinct transcriptional programs, all emerging as stable attractors of a single underlying GRN whose dynamics are predominantly explained by TF–DNA binding and protein–protein interactions from orthogonal molecular evidence.

**Highlights:**

- SETIA infers causal gene regulatory networks whose dynamics reproduce and generalize to held–out transcriptional profiles as stable attractors
- Genes occupy multiple discrete, reproducible expression states
- A ChIP–exo–derived protein–protein/DNA interaction network provides structural priors that ground the GRN in molecular mechanisms
- Molecular structural priors improve mechanistic interpretability while maintaining dynamical performance
- SETIA generalizes across bulk and single–cell data and scales to a semi–genome–scale regulatory network

## Introduction

Cells in multicellular organisms share an identical genome yet differentiate into diverse, specialized cell types through regulated gene–expression programs rather than genetic changes. This principle, known as genomic equivalence, was first demonstrated by nuclear transfer experiments showing that differentiated nuclei retain the potential to generate alternate cell types^1^. Differentiated cell states are stable yet reversible under specific circumstances, as illustrated by reprogramming somatic fibroblasts into induced pluripotent stem cells using four transcription factors (Oct3/4, Sox2, Klf4, and c-Myc), collectively known as the “Yamanaka factors”^2^. These findings highlight the plasticity of cell fate and the central role of transcription factors in controlling state transitions.

The high reproducibility of RNA–seq measurements across biological replicates suggests that distinct transcriptional profiles reflect stable, underlying cellular states rather than stochastic noise^3^. Rooted in Waddington’s epigenetic landscape and formalized in gene regulatory network theory, these stable states correspond to “attractor” states in the high–dimensional landscape of gene expression^4–8^, arising from network dynamics and corresponding to distinct cell fates^9,10^.

A comprehensive GRN should integrate molecular evidence, such as RNA, TF–DNA binding and protein–protein interactions, while capturing dynamics that maintain stable states and reproduce transcriptional responses to perturbations^11,12^. Such models should enable prediction of how cells reconfigure gene expression in response to genetic or environmental changes. However, achieving this integration remains a central challenge.

Existing approaches broadly fall into model–free and model–based methods^13^. Model–free approaches (e.g., ARACNe^14^, CLR^15^) infer associations from expression correlations but do not establish causality^16^. Model–based methods such as BGRMI^17^, BINGO^18^, EA^19^, and others^20–22^ explicitly model gene expression dynamics using ordinary differential equations (ODEs) or probabilistic state–space formulations. These models fit parameters to static or time–series expression data, yielding a parameterized dynamical model that specifies how regulators influence target genes.

Despite these advances, three limitations remain. First, most methods do not integrate multiple data modalities within a unified framework. Gene regulation is measured across orthogonal assays, including ChIP–seq^23^ and CUT&RUN^24^ to measure TF–DNA binding, PRO–seq^25,26^, CAGE–seq^27^, ChRO–seq^28^, and NET–seq^29^ to quantify nascent transcription and promoter activity, and RNA–seq to characterize steady–state gene expression. Yet these layers are rarely simultaneously reconciled during inference. Second, existing models typically assume only a few expression levels (e.g., off, low, high)^30^. It is not known whether gene expression occupies multiple discrete, reproducible states across conditions. Capturing these multi–level states provides stronger constraints on GRN structure. Third, modeling explicit dynamics scales poorly with network size, making inference computationally challenging^22,31^.

To address these challenges, we present SETIA, a machine–learning framework that integrates molecular evidence with dynamical simulation to infer predictive, mechanistically grounded GRNs. SETIA addresses all three limitations: it integrates orthogonal molecular data modalities, including ChIP–exo–derived TF–DNA binding and protein–protein colocalization as structural priors^32,33^; it models multi–level expression states (up to nine levels) rather than assuming binary on/off behavior; and it enables scalable inference by decomposing the GRN into per–gene sub–networks trained in parallel and then reassembled into an integrated GRN. This approach enables dynamical GRNs that reconcile orthogonal molecular evidence with multi–level transcriptional states while remaining computationally tractable.

## Results

### SETIA reconciles transcriptional dynamics with orthogonal regulatory evidence in a unified framework

SETIA was architected to model how transcription factors generate reproducible gene expression patterns across perturbations and conditions by integrating RNA–seq data with orthogonal regulatory evidence, including TF–gene promoter binding and TF–TF interactions. Rather than treating expression profiles as independent samples, SETIA requires that a single GRN reproduces all observed transcriptional profiles as stable states of its dynamical model (**Figure S1**), thereby enforcing a direct correspondence between network structure and observed transcriptional outcomes.

SETIA first establishes candidate regulatory relationships that specify an initial GRN structure (**Figure 1a**). TF–gene promoter binding data constrain potential regulatory edges, while TF–TF relationships inferred from protein–protein interaction data identify factors likely to act jointly on target genes. Together, these constraints define a molecularly informed yet flexible initial network.

**Figure 1.**
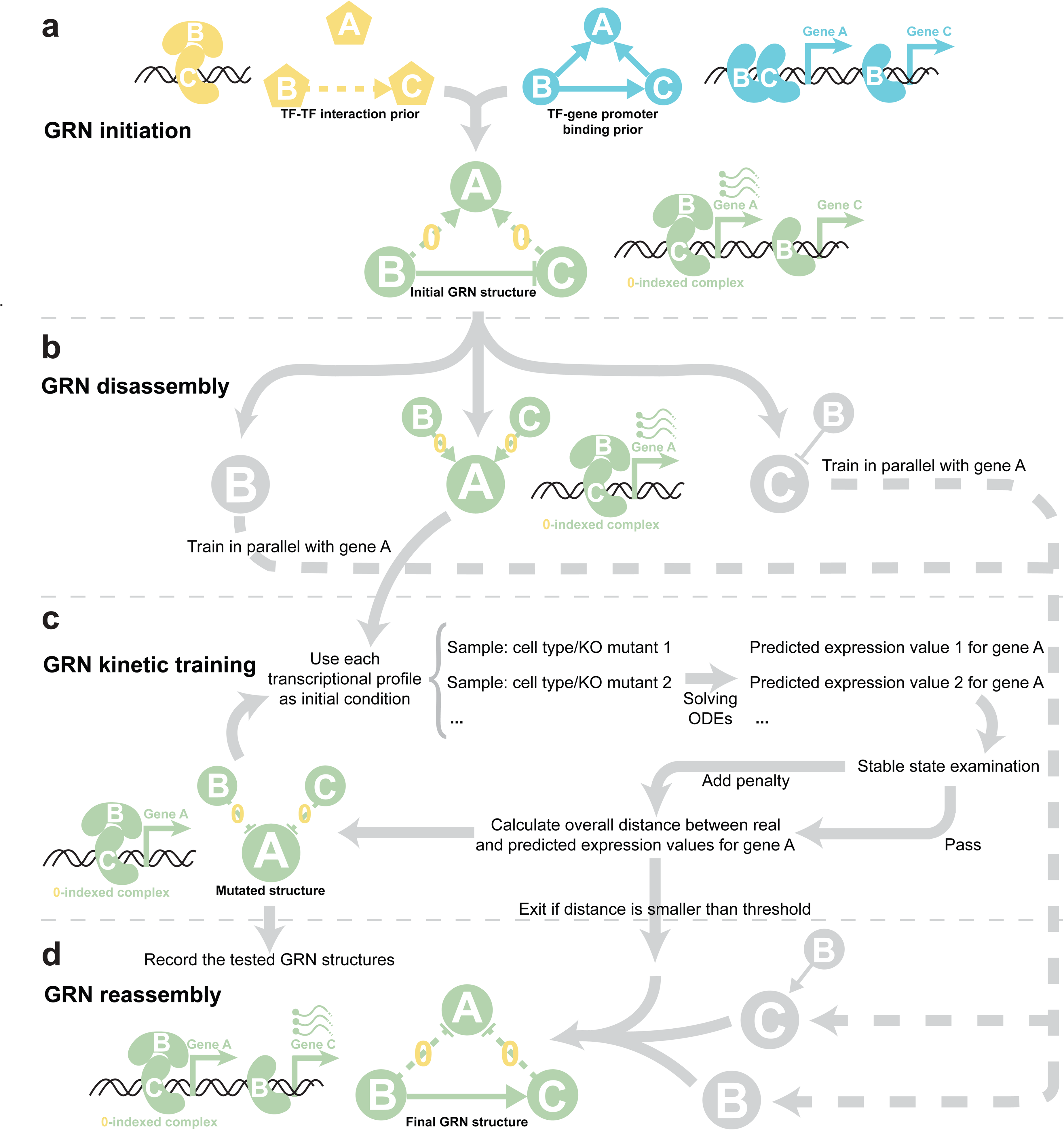
Schematic of the workflow of SETIA. (a) TF–gene promoter binding and protein–protein interaction compose the initial network structure. Solid edges denote TF–gene promoter binding specified by the TF–gene promoter binding prior and represent independent regulatory interactions, whereas dashed edges denote protein–protein interactions specified by the TF–TF interaction prior and represent cooperative regulatory relationships between TFs. (b) The GRN is decomposed into per–gene sub–networks that include each gene’s incoming edges. (c) With constrains from the TF–gene promoter binding and protein–protein interaction priors, sub–networks are trained in parallel to fit their discrete expression levels derived from the RNA–seq data. (d) Optimal sub–networks are reassembled into the final integrated GRN.

To enable scalability to large networks, SETIA’s design decomposes a GRN into gene–specific regulatory sub–networks (**Figure 1b**), each comprising a target gene and its candidate regulators. This unique aspect reduces computational cost by confining dynamical simulations to low–dimensional systems, allowing inference to be performed independently and in parallel across genes. Compared with joint ODE approaches, which scale exponentially with network size, this strategy substantially improves efficiency. For example, SETIA required ∼9 core–hours to infer a 9–gene GRN compared to ∼300 core–hours for an evolutionary–algorithm–based joint ODE approach^19^, representing roughly a 33–fold speedup. SETIA scaled to a 76–gene GRN in ∼35 core–hours, compared to an estimated 3450 billion core–hours with the alternative approach.

Each sub–network is trained to reproduce observed stable expression states (**Figure 1c**). Experimental profiles initialize the dynamical model and simulated steady–state expression is compared to observed values. Model parameters and structures are iteratively optimized to minimize discrepancies while satisfying molecular priors, retaining only solutions that consistently converge to the observed states.

Optimized sub–networks are then assembled into a single integrated GRN (**Figure 1d**) that reproduces input transcriptional profiles as stable states across conditions. This inferred GRN can be simulated from arbitrary initial states to evaluate convergence and generalization, enabling prediction of transcriptional responses to unseen perturbations, including TF knockouts and changes in regulatory interactions.

To facilitate exploration, we provide an interactive web–based GRN simulator (https://grn.cac.cornell.edu:5000/) for visualization, perturbation, and dynamical analysis. An overview of the GRN simulator website and usage guidelines is provided in **Figure S2**.

### Genes exist in multiple stable expression states

We modeled gene expression across conditions as a mixture of discrete stable states, consistent with the assumption that transcriptional profiles correspond to attractors of GRN dynamics. Constitutive genes are expected to exhibit a single unimodal distribution, whereas regulated genes are thought to exist in either an “on” or “off” state. However, if multiple regulators act on a particular gene, each making an “on/off” contribution, then in principle multiple stable transcriptional states might exist depending on regulator configuration.

To test this hypothesis in a controlled perturbation setting, we focused on a compact regulator set in yeast: the 76 sequence–specific transcription factors (ssTFs) defined by Rossi et al.^34^. We sequenced mRNA from wild–type and 36 single TF knockout (TFKO) *Saccharomyces cerevisiae* strains (37 conditions total; the full list of TFKOs is shown in **Figure 2c**; 36 rather than all 76 ssTFs were deleted because a fraction are essential for viability). Replicate samples clustered by genotype (**Figure S3**), supporting data reproducibility and genotype–specific transcriptional responses.

**Figure 2.**
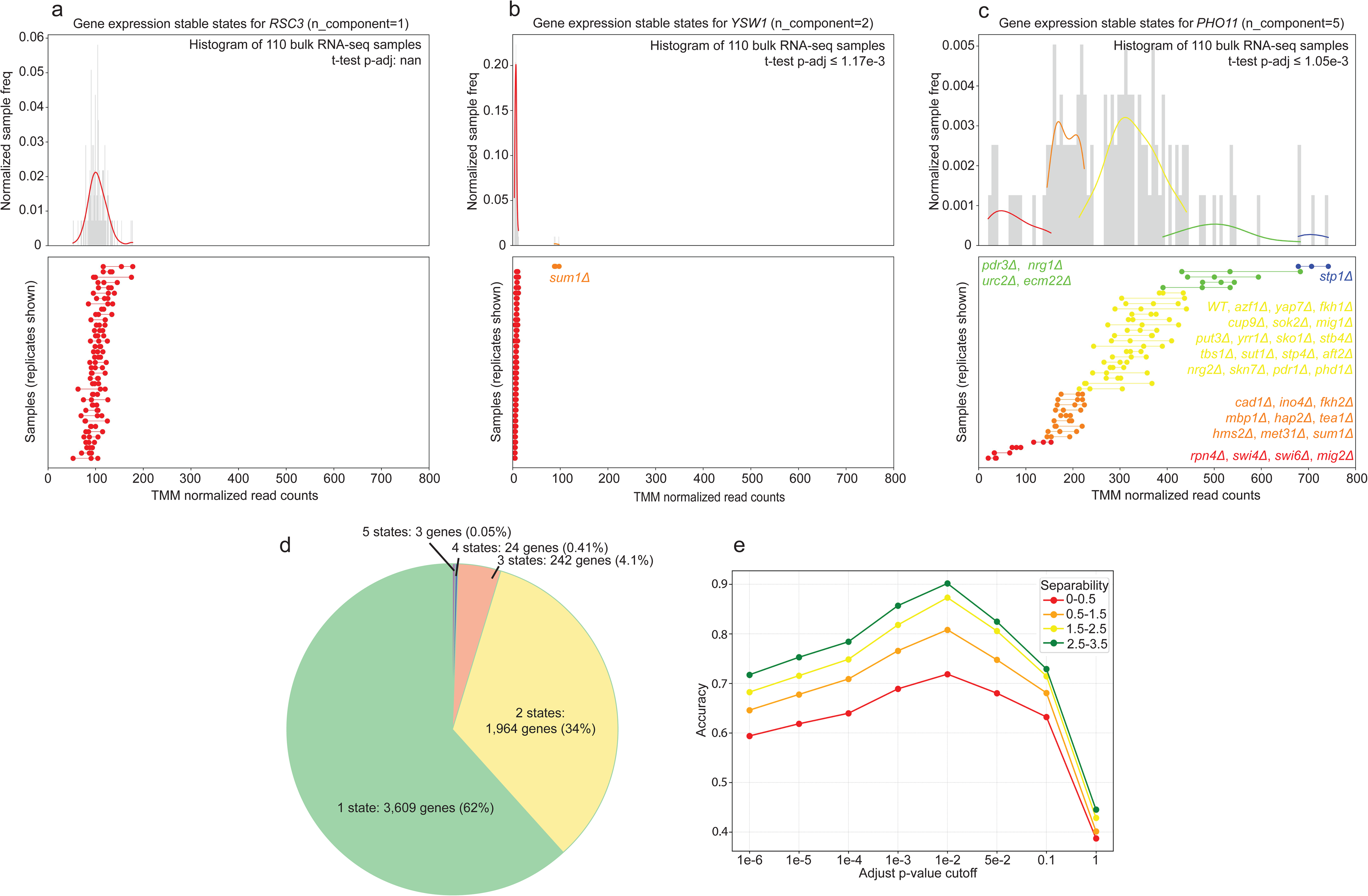
Gene expression exists in multiple stable states. (a–c) Three representative genes, *RSC3*, *YSW1*, and *PHO11*, illustrating distinct expression behaviors in response to TFKOs. The bottom panels show TMM–normalized read counts for each genotype, where each row corresponds to a different genotype and dots connected by a line represent biological replicates of the same genotype. The top panels summarize the same data as histograms of the samples, overlaid with kernel density estimates (KDEs). Genes with a single stable state display a unimodal distribution (a), whereas genes with multiple stable states exhibit clearly separated KDE modes (b, c). Welch’s t-tests were performed between all pairs of putative states with multiple–testing correction, and the largest adjusted p-value was reported in the top–right corner of each panel. (d) Proportion of genes exhibiting different numbers of stable expression states, as identified by **Algorithm 1** using TMM–normalized read counts from the biological replicates across the wild–type and 36 TFKO yeast strains. (e) Performance of the hybrid state–identification algorithm on an *in silico* benchmark with known ground truth. Synthetic gene expression values were generated from mixtures of Gaussian distributions with varying degrees of state separability. Classification accuracy is shown as a function of the adjusted p-value cutoff for different separability regimes, with optimal performance achieved near a cutoff of 0.01.

We next applied a replicate–aware state–calling algorithm (**Algorithm 1**) to infer discrete expression states from TMM–normalized read counts. Because the biological interpretation of inferred states depends on the reliability of this procedure, we evaluated its performance using an independent *in silico* benchmark with known ground truth (see STAR Methods). Synthetic datasets were generated by sampling replicate expression values from mixtures of Gaussian components, with each component representing a latent expression state and replicates constrained to arise from the same state. State separability was systematically varied by adjusting the distance between component means. Under these conditions, Algorithm 1 achieved ∼70–90% classification accuracy, with performance improving as separability increased (**Figure 2e**), indicating reliable recovery of multi–state expression patterns.

Having established the reliability of the state–calling procedure, we next examined its behavior across the 110 RNA–seq samples spanning the 37 WT and TFKO genotypes. Most genes were assigned a single unimodal component, whereas others exhibited multiple discrete expression states, resulting in multimodal mixtures. To illustrate these expression patterns, we present representative genes corresponding to unimodal expression, direct TF–mediated regulation, and potentially indirect regulation. *RSC3* displayed a single stable state (**Figure 2a**). *YSW1* shifted to a higher stable state upon *SUM1* deletion, consistent with its known Sum1–Hst1–Rfm1 chromatin repression pathway^35,36^ (**Figure 2b**). *PHO11* exhibited five distinct stable states, indicating reproducible multi–level regulation across perturbations rather than continuous variation (**Figure 2c**). Notably, in our ChIP–exo dataset only 2 of the 36 deleted ssTFs, Fkh1 and Sok2, were detected at the *PHO11* promoter, consistent with at least a subset of these stable states arising through indirect effects of TF deletion. For instance, deletion of *SWI4* or *SWI6* perturbs the G1/S transcriptional program controlled by the Swi4–Swi6 (SBF) complex and could alter *PHO11* expression through indirect transcriptional cascades or by enriching cells in particular cell–cycle phases, thereby changing bulk expression readouts of *PHO11*^37^.

Genome–wide, 2,233 genes (38.2%) exhibited more than one expression state, and 269 genes (4.6%) exhibited three or more (**Figure 2d**; **Supplementary Data 1**). These results indicate that gene expression in the WT and TFKO mutant strains is often organized into discrete, reproducible regimes rather than continuous variation.

### SETIA–inferred GRNs accurately reproduce stable transcriptional states

Having identified discrete and, in many cases, multi–state gene expression behaviors across the WT and TFKO yeast strains, we next asked what GRN structure is sufficient to generate these transcriptional profiles as stable states of a dynamical system. We treated the 37 WT/TFKO RNA–seq profiles as observed stable states and trained a GRN over the 76 ssTF–encoding genes using SETIA, such that its simulated dynamics recapitulate these transcriptional profiles as stable states, including genes exhibiting multiple distinct expression modes (**Figure 3a**; **Supplementary Data 2**).

**Figure 3.**
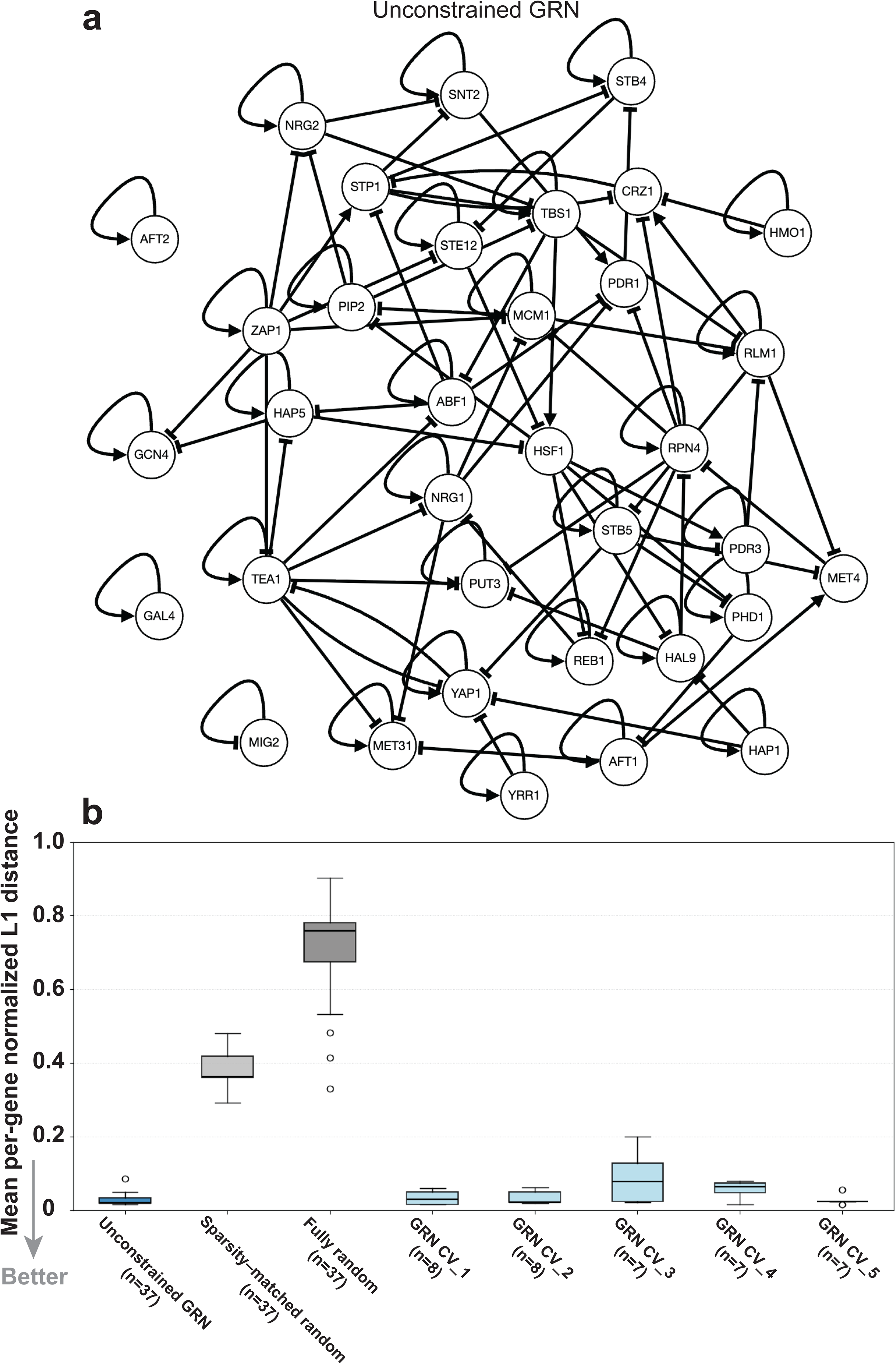
Unconstrained GRN inference and dynamical validation through stable–state reproduction. (a) Gene regulatory network inferred by SETIA without imposing structural priors. Genes exhibiting a single stable expression state across the bulk RNA–seq dataset are omitted for clarity. Edges represent effective regulatory influences inferred from transcriptional dynamics and could reflect indirect or composite effects rather than direct molecular interactions. (b) Dynamical performance of the inferred GRN (blue) compared with two random controls, a fully random (dark gray) and a sparsity–matched random GRN (light gray). Each transcriptional profile (37 WT and TFKO conditions) was used as an initial condition, and the system was simulated forward in time to assess whether it is recovered as a stable state. The inferred GRN preserved nearly all profiles as stable states, as indicated by near–zero distances between initial and final states, whereas both random controls failed to do so. Five–fold cross–validation results (CV_1–CV_5) are also shown, in which transcriptional profiles were partitioned into non–overlapping subsets and held–out profiles were tested for recovery as stable states. The number of transcriptional profiles evaluated for each GRN is indicated by n on the x–axis. Distances were quantified using the mean per–gene normalized L1 metric.

We then evaluated the inferred GRN by testing whether it reproduced all 37 profiles as stable states. For each profile, we initialized the system to the observed 76–gene expression vector and simulated forward in time by numerically integrating the ODEs defined by the inferred GRN (see STAR Methods). If the initialized profile is a stable state of the GRN dynamics, the trajectory remains near its starting point; otherwise, it will relax to a different state, producing a large distance between the initial and final states. We quantified this deviation using the mean per–gene normalized L1 distance between the initial and final expression vectors (see STAR Methods). We then compared the SETIA–inferred GRN to two randomized baseline networks. One was a fully random GRN in which one of three edges (activation, inhibition, or no edge) was assigned independently to each ordered TF–target pair. The other was a network–sparsity–matched random GRN obtained by shuffling the edges of the inferred GRN. This edge–shuffling procedure preserves the total number of regulatory edges, and therefore the overall network sparsity, while randomizing their source and target assignments. The SETIA–inferred GRN recovered all 37 transcriptional profiles as stable states, whereas neither randomized control reproduced the profiles as stable attractors (**Figure 3b**). This result demonstrated that the inferred GRN can encode the causal, dynamical relationships among the selected TFs, so that a TF’s stable–state expression level is determined by the expression of its regulators, and the network dynamics preserve those relative levels across all input cell–type–specific transcriptional profiles.

To evaluate the predictive capacity of unconstrained GRNs, we next asked whether a fully unconstrained GRN can generalize beyond the transcriptional profiles used for inference. Conventional GRN validation typically relies on comparing predicted edges to chromatin accessibility, TF–DNA binding maps, or literature–derived regulatory networks^38,39^. Although such benchmarks assess whether individual interactions are plausible, they do not test whether an inferred GRN can reproduce transcriptional states as dynamical outcomes under perturbation, a criterion that has been proposed as a key principle for evaluating GRN models^40^. Because SETIA infers an explicit dynamical model, we can evaluate whether simulated GRN dynamics recover measured transcriptional profiles and generalize to those not used during inference. We therefore performed a five–fold cross–validation in which the 37 TFKO and WT transcriptional profiles were randomly partitioned into five non–overlapping subsets, leaving out one–fifth of the transcriptomic profiles in each fold without replacement. For each fold, a GRN was inferred from the remaining profiles, and the system was simulated forward from each held–out initial condition to test whether the excluded profiles emerged as stable states of the inferred dynamical system. Across all folds, the unconstrained GRNs consistently recovered the held–out transcriptional profiles as stable states (**Figure 3b**). These results indicate that unconstrained edges capture biologically meaningful regulatory influences that enable robust dynamical generalization, allowing the model to recover unseen transcriptional stable states arising from the same underlying regulatory program.

Although the inferred GRN reproduces transcriptional dynamics, it was optimized without requiring agreement with orthogonal molecular evidence. Consequently, its edges represent effective influences, including both direct TF–gene regulation and indirect dependencies arising from unmodeled factors or cellular processes. We next incorporated independent molecular evidence to constrain the inferred GRN toward measured direct regulatory interactions.

### A promoter–architecture–aware TF–gene binding network robustly grounds GRN inference

To improve the molecular interpretability of inferred regulatory edges, we incorporated two complementary layers of structural prior evidence into SETIA: TF–gene promoter binding and protein–protein colocalization. As a first layer, we constructed a TF–gene promoter binding network by integrating site–specific TF–DNA binding data and annotated nucleosome–depleted or free regions (NDRs/NFRs) from Rossi et al.^34^. NDRs/NFRs within promoter regions were treated as candidate regulatory sites at which transcription factors bind to control target gene expression. In the densely packed *S. cerevisiae* genome, promoter NDRs/NFRs frequently reside within bidirectional intergenic regions shared by head–to–head gene pairs, and promoter boundaries can overlap with adjacent transcriptional units in head–to–tail configurations. We therefore incorporated gene arrangement and insulator binding (Abf1, Rap1, or Reb1) to more realistically map ChIP–exo peaks in promoter NDRs/NFRs to TF–target gene regulatory interactions. For tandem head–to–tail gene configurations, a TF–target gene interaction was assigned when a ChIP–exo peak was detected within the target gene’s promoter NDR/NFR (**Figure S4a**). For divergent head–to–head gene pairs, we evaluated the distance between their open reading frames to assess potential co–regulation by the shared intergenic region. When the intergenic distance was less than 300 bp, the shared promoter was assumed to regulate both genes. For distances between 300 bp and 700 bp, co–regulation was assumed unless one or more insulator proteins were bound within 30% of the intergenic region. When the distance exceeded 700 bp, the two genes were not considered to be co–regulated (**Figure S4b**).

We first parsed adjacent gene orientations genome–wide and identified 3,161 tandem (head–to–tail) genes and 2,434 divergent (head–to–head) genes based solely on the relative orientations of neighboring ORFs. We then applied the promoter–sharing criteria, intergenic distance thresholds and insulator binding as described above, to define a subset of divergent gene pairs with a shared bidirectional promoter. Using the same underlying data as Rossi et al. but an independent analytical approach, we identified 811 such divergent head–to–head protein–coding gene pairs, including all 762 pairs reported by Rossi et al. (**Figure S4c**). This concordance indicates that our approach recapitulates the established gene arrangement annotations reported by Rossi et al. while extending the set of identified divergent gene pairs under the same criteria.

Using our promoter definitions and ChIP–exo peak assignments, we constructed a TF–gene promoter binding network containing 5,864 nodes and 6,955 edges, provided as **Supplementary Data 3**. Here, nodes represent both structural genes and TF–encoding genes, and an edge denotes TF occupancy at a target–gene promoter, evidenced by a ChIP–exo peak within the defined promoter region. This particular criterion does not require the presence of the TF’s cognate DNA–binding site at the bound location. Accordingly, TF occupancy at a promoter may reflect either direct site–specific binding or indirect association mediated by interactions with other promoter–bound factors. Of the 6,955 edges in our network, 5,783 overlapped with the TF–DNA binding network reported by Rossi et al. in their **Supplementary Data 2**, indicating strong overall agreement between the two approaches. The remaining 1,912 edges present in the Rossi et al. network (7,695 total) but absent from ours primarily reflected cases in which no ChIP–exo peak fell within our promoter definitions, or instances in which a peak was assigned to the opposite gene in a head–to–head pair under our promoter assignment criteria. Conversely, we identified 1,172 edges unique to our network, each supported by a ChIP–exo peak within the corresponding promoter region. Together, these comparisons show that our network largely recapitulates previously reported TF–DNA interactions while adding high–confidence promoter–associated TF–gene edges directly supported by ChIP–exo evidence (**Figure S4d**).

### A genome–bound protein–protein interaction network further grounds GRN inference

While the TF–DNA binding network identifies TF–target gene associations, its inference from bulk ChIP–exo data does not distinguish whether TFs bind a promoter simultaneously or independently across the cell population. To capture combinatorial regulation arising from cooperative versus independent TF activity, we inferred a ChIP–exo–derived protein–protein colocalization network as a complementary structural prior for GRN inference. In ChIP–exo, two TFs that interact on chromatin will crosslink to each other and to DNA. Subsequent immunoprecipitation of each TF and exonuclease digestion at single–bp resolution produces essentially identical genomic peak locations for the interacting TFs and non–identical neighboring peaks for non–interacting TFs^34^. We therefore analyzed the ChIP–exo binding patterns of ∼400 genome regulatory proteins, centering on peaks identified by ChExMix^41^ and stratifying sites based on whether the ChExMix peak contained the cognate DNA motif of the assayed factor.

We quantified similarity between ChIP–exo composite profiles using two complementary metrics. The Jensen–Shannon divergence distance (JSDD) measures global dissimilarity between normalized crosslinking distributions of two factors and is sensitive to differences in peak position and shape (**Figure 4a**). Complementarily, the K ratio quantifies relative signal scaling at corresponding peak regions independent of positional differences. It represents the prominence–weighted proportionality constant relating mean crosslinking intensities between two factors across shared peak regions, capturing amplification or attenuation of signal at aligned peaks. These two metrics capture different aspects of ChIP–exo profile similarity: low JSDD indicates preservation of overall crosslinking architecture, whereas high K values indicate selective amplification of ChIP–exo signal. We then constructed distributions of JSDD and K ratio values for all factors relative to each mapped genome regulatory protein. We identified candidate interactors whose genome–wide composite ChIP–exo profiles were most similar to each other by analyzing the second derivative of the JSDD between composite profiles (**Figure 4b**). As the similarity threshold was relaxed, the JSDD values reached a plateau in which a large majority of factors exhibited nearly identical JSDD values. Because most of these proteins are not expected to interact with the queried protein, this plateau defines a natural background similarity level. In parallel, we applied an analogous second–derivative–based criterion to the K ratio distribution to identify factors exhibiting queried protein–specific, peak–localized enrichment. Factors meeting these criteria were designated as potential interacting cofactors for the queried protein. Applying this query to Spt15 (TATA binding protein) as an example, the five factors with the lowest JSDD values were Sua7, Tfa1, Tfa2, Kin28, and Cet1. In addition, the top five factors ranked by the K ratio were Kin28, Bdp1, Tfa2, Sua7, and Brf1. These factors are core Pol II initiation components such as Sua7 (TFIIB), Tfa1–Tfa2 (TFIIE), and Kin28 (TFIIH kinase module)^42–44^, as well as TBP–associated Pol III initiation factors Brf1 and Bdp1 (TFIIIB)^45,46^, and the Pol II–coupled mRNA capping enzyme Cet1^47^, providing independent biological support for the JSDD–K framework.

**Figure 4.**
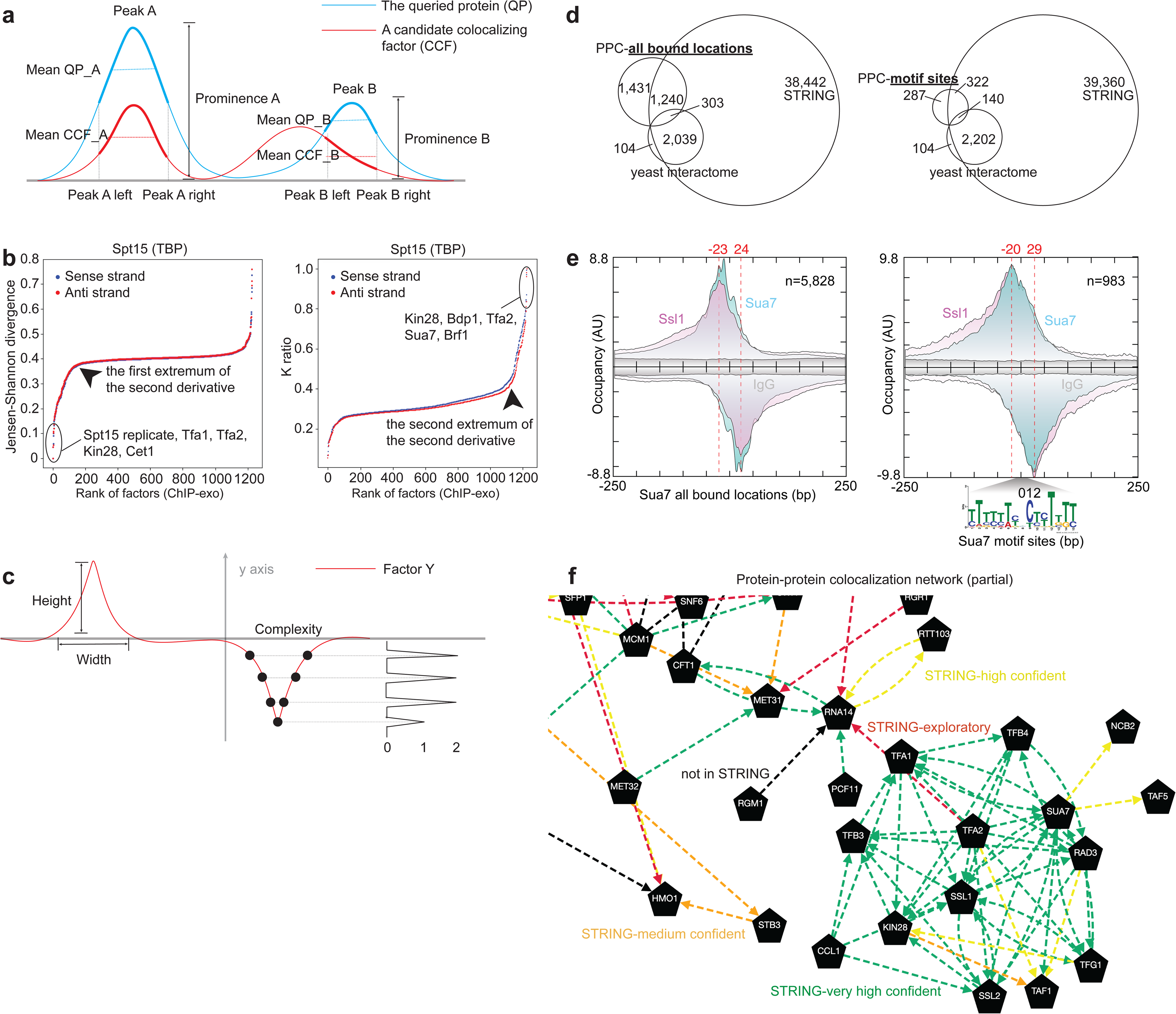
In addition to protein–DNA interactions, ChIP–exo defines protein–protein interactions across the genome. (a) The K ratio is a scalar that minimizes the prominence–weighted Euclidean base–pair distance between the average ChIP–exo signal within peaks identified for the queried protein and the corresponding regions in a candidate colocalizing factor. (b) Distributions of Jensen–Shannon divergence distance and K ratio comparing the ChIP–exo composite profile of the queried protein (Spt15) to those of all 1,229 candidate colocalizing factors. Potential cofactors with similar ChIP–exo profiles were identified by locating the first extremum in the second derivative of each distribution, corresponding to a natural separation between closely matching and dissimilar profiles. The top five matching factors are shown. (c) The secondary sorting was based on the peak height, width and complexity, or read depth, of the ChIP–exo profiles of candidate colocalizing factors. (d) Venn diagram illustrating the overlap of edges among our PPC networks, the yeast protein interactome, and the STRING database. (e) Overlapping ChIP–exo composite profiles of Sua7 and Ssl1 on Sua7 bound sites, stratified by the presence or absence of a cognate motif. (f) Example PPC network inferred from genome–wide colocalization, visualized using our GRN simulator website. Factors are represented as pentagon–shaped nodes, and edges are color–coded by STRING confidence levels.

To further refine the set of potential cofactors for the queried protein, we applied a secondary ranking based on a confidence score that captures the reliability of the composite ChIP–exo profiles. While the JSDD quantifies overall similarity in profile shape, it does not assess whether the features within a composite profile are supported by sufficient signal, particularly when low read depth produces jagged or fragmented peak patterns. As a result, apparent similarities in composite profiles may reflect noise rather than robust binding features. We therefore incorporated peak–level metrics that evaluate the quality of these features, including peak prominence, width, and complexity (see STAR Methods) (**Figure 4c**). Peaks with greater prominence, sufficient width, and higher complexity were assigned higher confidence, ensuring that selected cofactors not only share similar composite profile shapes with the queried protein but also exhibit reliable and biologically meaningful binding patterns.

Having defined a set of high–confidence protein–protein colocalization relationships based on composite ChIP–exo profiles, we next examined how these inferred interactions relate to existing protein–protein interaction resources. We constructed two protein–protein colocalization (PPC) networks, one based on all bound locations and one based on motif–bound sites (provided as **Supplementary Data 4** and **Supplementary Data 5**), and compared them with the yeast protein interactome from Michaelis et al.^48^ and the protein–protein interaction database named STRING^49^ (**Figure 4d**). From STRING, we included interactions classified as “very high confidence”, “high confidence”, “medium confidence”, and “exploratory”. In the all–bound–sites PPC network, 1,431 edges (48.1%) were unique to our PPC network, whereas 1,543 edges (51.9%) overlapped with STRING interactions, of which 303 (10.2%) were also present in the yeast interactome. For the motif–based PPC network, similar proportions were observed: 287 edges (38.3%) were unique to the PPC network, 462 edges (61.7%) overlapped with STRING, and 140 edges (18.7%) were additionally supported by the yeast interactome. The unique edges identified in our PPC networks suggest potential protein–protein interactions that may occur specifically *in vivo* or reflect close spatial proximity without requiring direct physical interaction.

Across both PPC networks, we identified 2,671 factor pairs with overlapping ChIP–exo composite profiles, including well–established pairs such as Sua7 and Ssl1 (**Figure 4e**). The extent of genome–wide colocalization was encoded as edge opacity. Edge colors were used to denote STRING confidence categories. The PPC networks and associated ChIP–exo composite profiles can be interactively explored using our GRN simulator website (**Figure 4f**). The union of the two PPC networks and STRING interactions classified as “very high confidence” was used for downstream GRN inference.

Together, these results demonstrate that PPC networks capture chromatin–context–specific and potentially directional TF associations that extend beyond known stable protein–protein interactions. By identifying TFs that co–occupy and are therefore likely to act jointly at regulatory elements, the PPC network provides structural priors that bias SETIA’s inference toward combinatorial regulation by favoring models in which co–localizing TFs synergistically regulate shared targets. This reduces ambiguity in GRN inference and enables dynamical models that reflect coordinated transcriptional control.

To investigate colocalization edges absent from existing protein–protein interaction databases, we examined representative cases using ChIP–exo composite profiles. This revealed instances of asymmetric, or directional, colocalization, in which the association of one factor depends on another. For instance, the ssTF Skn7 binds 104 genomic loci, whereas Sok2 binds 388. Composite profiles show strong colocalization of Sok2 at sites bound by Skn7, but substantially weaker overlap of Skn7 at Sok2–bound sites, indicating directional association rather than mutual binding (**Figure 5a** and **Figure 5b**). Quantitatively, Sok2 colocalizes with 66 of the 104 Skn7–bound sites, while the remaining 38 Skn7 sites lack a Sok2–like ChIP–exo profile (**Figure 5c** and **Figure 5d**). Together, these observations support a model in which Sok2 associates with Skn7 at a specific subset of regulatory loci in a context–dependent manner, consistent with *in vivo* colocalization at chromatin rather than formation of a stable, bidirectional protein–protein complex. Such directional and conditional TF associations are captured by the PPC network and incorporated into SETIA as structural priors that bias inference toward models in which these TFs act jointly or conditionally on shared targets. This enables the GRN to represent context–dependent and potentially synergistic regulatory relationships.

**Figure 5.**
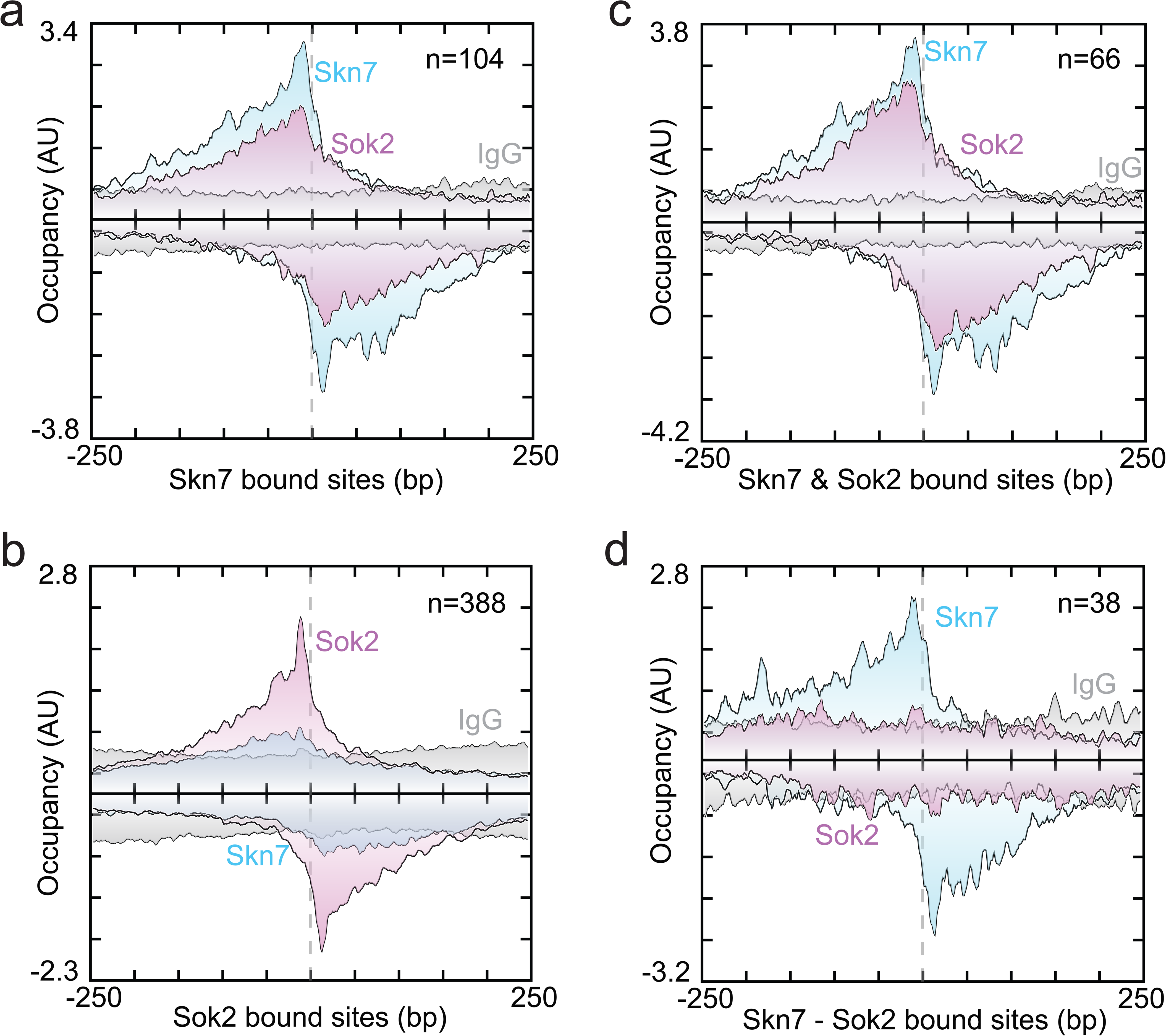
Genome–wide site–specific protein–protein interactions demonstrated by Skn7 and Sok2. (a) Composite ChIP–exo profiles of Skn7 and Sok2 centered on the 104 binding sites occupied by Skn7. (b) Composite ChIP–exo profiles of Skn7 and Sok2 centered on the 388 binding sites occupied by Sok2. (c) Composite ChIP–exo profiles of Skn7 and Sok2 centered on the 66 binding sites co–occupied by both factors. (d) Composite ChIP–exo profiles of Skn7 and Sok2 centered on the 38 Skn7–bound sites that do not show Sok2 occupancy.

### Structural priors improve mechanistic interpretability while constraining dynamical stable state generalization

Although an unconstrained GRN can reproduce observed transcriptional profiles as stable states, this does not uniquely determine the underlying network molecular structure, as indirect or unmodeled effects can be absorbed into effective regulatory edges. This limitation is not specific to SETIA but reflects a general challenge in GRN inference approaches that rely primarily on transcriptional data. To incorporate mechanistic constraints, we inferred GRNs under structural prior information derived from TF–gene promoter binding and protein–protein colocalization or interaction, aiming to reduce reliance on uninterpretable indirect effects.

We inferred four GRN variants that differed only in the strength and form of structural constraints imposed during inference (**Figure 6a**). GRN A, also shown in **Figure 3a**, was inferred without any structural priors and therefore allowed all regulatory edges supported by the dynamical fitting procedure. GRN B incorporated TF–gene promoter binding together with gene–gene expression correlation, such that an edge was permitted if there was either a TF–gene promoter binding event or evidence of co–expression between the two genes (**Supplementary Data 6**). GRN C restricted edges exclusively to those supported by TF–gene promoter binding events in ChIP–exo data. Co–binding of multiple TFs provides evidence of co–regulation. For example, Stp1, Aft2, and Yap1 co–localize and converge on *AFT1* in GRN C, forming a regulatory module consistent with synergistic regulation (**Figure 6b**; **Supplementary Data 7**). GRN D imposed the most stringent constraint by further requiring the presence of a cognate DNA motif at each TF–bound site, thereby enforcing both physical binding and sequence specificity (**Supplementary Data 8**).

**Figure 6.**
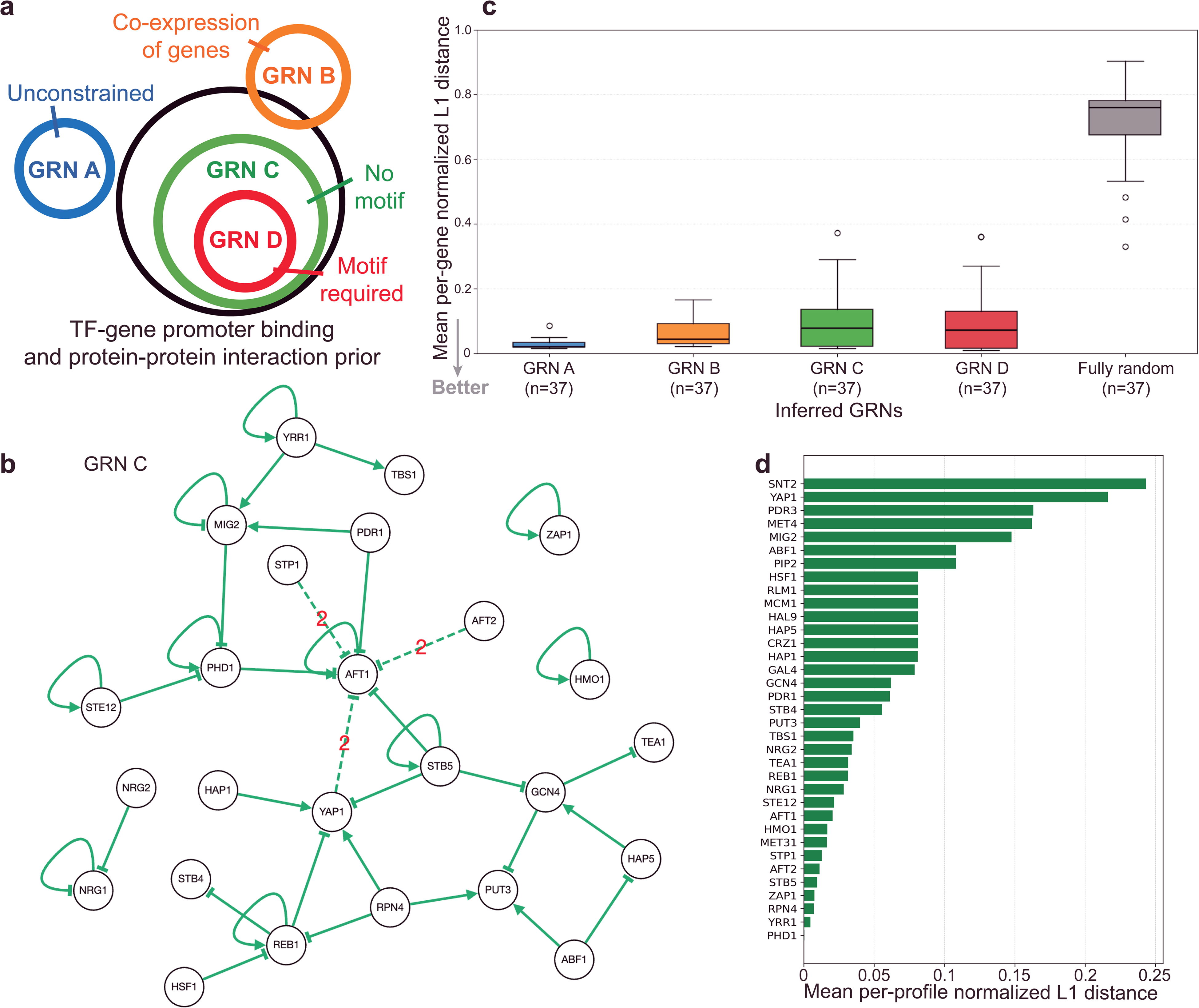
Influence of structural priors on inferred GRN dynamical performance. (a) Schematic overview of the four GRN inference regimes, A–D, with increasingly strict structural constraint. (b) Network architecture of GRN C, in which all regulatory edges are supported by the TF–gene promoter binding prior; edges are shown in green to indicate agreement with the prior. Genes exhibiting a single expression state across the dataset, as well as isolated genes lacking regulatory connections, are omitted for clarity. (c) Dynamical performance of GRNs A–D compared with a random GRN control in reproducing 37 transcriptional profiles as stable states, quantified using the mean per–gene normalized L1 distance. The number of transcriptional profiles evaluated for each GRN is indicated by n on the x–axis. (d) Gene–wise prediction errors for the TF–gene promoter binding constrained GRN (GRN C), revealing that large errors are concentrated in a small subset of genes whose observed multi–state expression behavior is incompatible with strict TF–gene promoter binding constraints, highlighting conflicts between hard structural priors and transcriptional data.

All four GRNs performed well on reproducing the 37 transcriptional profiles, with median mean per–gene normalized L1 distances below 0.1, compared to 0.7 for a random GRN (**Figure 6c**). Thus, a substantial fraction of the observed transcriptional variation among TFKO strains is explainable by direct ssTF–promoter regulation. A modest decline in dynamical performance was observed as GRN priors were strengthened (A→D), which reflects reduced flexibility to capture indirect regulatory effects outside the modeled TF set. This modest performance cost indicates that site–specific TF binding at promoters is the most dominant aspect of transcriptional regulation, validating the molecular basis of the network as defined by GRN D.

To understand the source of the reduced agreement observed under strict structural constraints, we performed a gene–wise error analysis using the mean per–profile normalized L1 distance, a metric appropriate for comparing gene–level prediction errors averaged across transcriptional profiles. This analysis revealed that prediction errors were concentrated in a small subset of genes, with the largest per–gene errors observed for *SNT2*, *YAP1*, *PDR3*, and *MET4* (**Figure 6d**).

For *SNT2* and *MET4*, the RNA–seq data exhibit multiple discrete expression states across samples, yet none of the differentially expressed ssTFs among the 76 modeled factors bind their promoters in our TF–gene promoter binding prior. Consequently, under the strict TF–gene promoter binding constraints embodied by GRNs C and D, these loci were effectively treated as unregulated single–state genes, an assumption incompatible with their observed multi–state expression and therefore a direct explanation for the poor fits. For *YAP1* and *PDR3*, multiple TFs in the prior are supported as binding their promoters. However, both genes occupy a larger number of discrete stable expression states than most other genes, increasing the dynamical complexity required to reproduce their behavior and making them correspondingly more difficult to fit under strict structural constraints. Together, these observations indicate that the largest prediction errors arise specifically from regulatory behaviors not represented in the hard structural prior, rather than from limitations of the dynamical inference itself, arguing for treating structural priors as soft constraints when the goal is to faithfully reproduce transcriptional profiles as dynamical stable states.

Overall, these results demonstrate that multiple GRN architectures can reproduce the same transcriptional stable states, while differing in the aspects of the system they optimize. Unconstrained GRNs generalize effectively to unseen stable transcriptional states by absorbing indirect regulatory influences into effective edges, whereas structurally constrained GRNs prioritize molecular interpretability and alignment with orthogonal molecular evidence. Accordingly, the appropriate degree of prior stringency should be chosen to match the analytical goal, whether maximizing predictive performance and stable–state generalization or inferring a mechanistically interpretable, evidence–grounded GRN structure.

### SETIA generalizes to single–cell–derived expression data and yields selective regulatory predictions

Having characterized how varying degrees of structural prior stringency shape GRN architecture and dynamical performance, we next asked whether SETIA’s core inference principles generalize beyond the bulk yeast perturbation data used above. Specifically, we tested whether dynamical GRN inference can extract consistent regulatory signal from single–cell–derived expression data across broader condition contexts without imposing structural priors, reserving independent binding evidence for downstream benchmarking. To this end, we evaluated SETIA on data derived from single–cell RNA–seq in *Saccharomyces cerevisiae* comprising wild–type and 12 transcription factor deletion strains profiled under 11 growth conditions^59^. Cells sharing the same genotype and growth condition were aggregated into pseudobulk profiles, which were then used as input to SETIA. As in the bulk mRNA–seq analyses, GRN inference was restricted to the 76 ssTF–encoding genes and performed without imposing TF–gene promoter binding or protein–protein interaction priors.

Notably, the single–cell–derived pseudobulk profiles recapitulated discrete gene expression states analogous to those observed in bulk RNA–seq data (**Figure S6a**). Globally, there was substantial concordance between the two datasets in genes classified as single–state (**Figure S6d**). Inspection of individual genes revealed both unimodal expression distributions (e.g., *ARP2*) and genes occupying multiple discrete expression states across conditions and cell types (e.g., *POR1*; **Figure S6b** and **Figure S6c**). These results confirm that the discrete attractor structure observed in bulk RNA–seq data extends to single–cell–derived profiles, supporting the use of this dataset for GRN inference benchmarking.

For benchmarking, we compared SETIA to Inferelator 3.0, a regression–based GRN inference framework designed to learn shared and condition–specific regulatory interactions^38^. Inferelator was applied to the same pseudobulk single–cell dataset in a prior–free setting, and regulatory edges were selected using the confidence cutoff automatically determined by Inferelator for this dataset (confidence score ≥0.3788). Because our analysis was restricted to 76 ssTFs, the inference problem was limited to 76 × 76 = 5,776 possible directed regulator–target edges. Under this restriction, direct comparison against curated yeast regulatory resources becomes problematic. When existing gold standard networks^38,39^ are filtered to include only interactions among these 76 TFs, only 14 annotated edges remain. This extreme sparsity renders standard benchmarking metrics largely uninformative, as most inferred edges cannot be evaluated against known interactions. To enable a more meaningful evaluation, we therefore defined a permissive reference set as the union of the curated gold standard and the TF–gene promoter binding network constructed in this study. Within the resulting 5,776–edge search space, 168 regulator–target edges supported by either curated regulatory evidence or direct TF–gene promoter binding were treated as positives, while the remaining 5,608 unsupported edges were treated as negatives for evaluation.

Using the 5,776–edge search space and the permissive 168–edge reference, Inferelator predicted 349 regulatory edges, of which 17 matched the reference set. In contrast, SETIA predicted 76 edges, with 7 reference–supported matches. Thus, Inferelator recovered a larger absolute number of reference interactions, whereas SETIA produced a smaller and more selective set of predictions. Given the highly imbalanced evaluation space, with positive edges comprising less than 3% of all possible interactions, performance was assessed using precision, recall, F1 score, and Matthews correlation coefficient (MCC), which more appropriately capture the tradeoff between selectivity and coverage. Under these metrics, SETIA achieved higher precision and a higher MCC. Inferelator attained higher recall and F1 score (**Figure S5a**), which is consistent with SETIA prioritizing reliability over coverage and Inferelator favoring broader exploratory inference. Importantly, these edge–level metrics assess only static overlap with a reference network and do not evaluate whether inferred networks can explain transcriptional states as emergent properties of regulatory dynamics. SETIA infers an explicit dynamical GRN and can therefore be evaluated by its ability to reproduce observed transcriptional profiles as stable states and to generalize to unseen profiles under the same condition. In contrast, Inferelator infers static regulatory associations without an explicit dynamical framework for testing stability or attractor behavior.

Although SETIA and Inferelator exhibit comparable performance in recovering regulatory interactions within the ssTF network, the two methods differ fundamentally in their computational design and intended scope. Inferelator is optimized for genome–wide GRN inference using regression–based algorithms, whereas SETIA explicitly models GRNs as systems of ordinary differential equations whose stable states represent observed transcriptional profiles. Extending SETIA directly to genome–scale inference will require numerical integration of large systems of ordinary differential equations and remains computationally challenging. To address this scalability gap while preserving an explicit dynamical interpretation, we adopted a hybrid semi–genome–scale modeling strategy, described in the following section.

### A semi–genome–scale hybrid strategy extends SETIA to genome–wide transcriptional outputs

To extend SETIA beyond the 76 ssTF network while preserving dynamical interpretability, we adopted a hybrid semi–genome–scale strategy in which the ssTF GRN serves as a dynamical core, and non–TF genes are connected as downstream targets through direct regulatory edges supported by TF–DNA binding evidence. In the resulting semi–genome–scale GRN, edges from ssTFs to non–TF genes were permitted only when supported by TF–DNA binding, and non–TF genes were modeled as outputs that do not feed back into the core network. This construction preserves the ability of the ssTF core to reproduce its transcriptional profiles as stable states, while extending the model to genome–wide transcriptional outputs.

We evaluated this semi–genome–scale GRN (**Figure S5b**) on 714 genes (32% of the 2,233 differentially expressed genes) that both exhibited more than one discrete expression state in the bulk RNA–seq data and were bound by at least one of the 76 ssTFs. For each gene, we computed the mean per–profile normalized L1 distance between observed and model–predicted expression levels and summarized these errors across genes (**Figure S5c**). The resulting distribution has a mean L1 distance of 0.185, indicating that predicted expression levels deviate, on average, by less than 20% of each gene’s normalized expression range across profiles. These results demonstrate that the semi–genome–scale GRN provides an accurate approximation of transcriptional outputs for non–TF genes under direct ssTF regulation, and that regulatory relationships learned during TF–centered inference generalize to downstream targets, supporting the extension of SETIA to broader, semi–genome–scale transcriptional modeling.

The full semi–genome–scale GRN is provided in **Supplementary Data 9** and can be interactively explored using our GRN simulator website.

## Discussion

A foundational premise of SETIA is the classic attractor view of gene regulation, originally mathematically articulated by Kauffman, in which cell types correspond to stable attractors of a GRN and perturbations drive transitions between them^50,51^. Under this framework, a GRN admits a finite number of dynamical stable states, implying that the expression of a gene should occupy a discrete set of stable expression regimes across conditions^7^. Consistent with this prediction, we found that many genes in *S. cerevisiae* exhibit multiple discrete expression states in both bulk and single–cell RNA–seq datasets, including a substantial subset with more than three stable modes. In the single–cell RNA–seq dataset, genes such as *ARP2* show unimodal expression, whereas *POR1* exhibits multiple discrete expression states. We also identified a substantial cohort of genes with effectively single–state behavior across conditions, with considerable overlap between bulk and single–cell datasets. This overlap suggests a robust set of genes whose expression remains effectively invariant across the sampled perturbations and conditions. Together, these observations support a model in which gene expression is organized into discrete attractor states, with a stable core of genes and a subset that transitions between states in response to genetic and environmental perturbations.

Building on this attractor–based view of transcriptional regulation, our results clarify how GRNs should be inferred when the goal is to reproduce observed transcriptional stable states while retaining mechanistic meaning. Existing approaches have addressed this challenge in different ways, either by prioritizing predictive performance from expression data alone^52,53^ or by imposing structural priors derived from TF binding, chromatin accessibility, or protein–protein interactions to improve interpretability^11,54^. However, these methods typically do not require inferred networks to reproduce observed transcriptional profiles as stable dynamical states, nor do they explicitly test whether predictive generalization and mechanistic structure can be reconciled within a single GRN framework. SETIA directly addresses this gap by inferring GRNs whose dynamics are constrained to reproduce transcriptional stable states, while allowing structural priors to be incorporated flexibly to balance predictive accuracy and biological interpretability.

At one extreme, SETIA can be run without structural priors, allowing the inferred GRN to focus solely on the dynamical requirement that the RNA–seq profiles correspond to fixed–point attractors. In this structurally unconstrained mode, the inferred GRN recapitulated essentially all 37 WT and TFKO transcriptional profiles as stable states. Although unconstrained GRNs may include effective edges that absorb indirect regulatory influences, this regime represents a valid and complementary inference mode in which relaxing structural constraints is appropriate, particularly because TF occupancy and transcriptional output are often measured in different biological contexts. For example, TF occupancy can be modulated by post–translational modifications that alter binding affinity, and many TFs can be poised at promoters such that binding is permissive while regulatory output remains stimulus–dependent. In such cases, relaxing structural priors allows SETIA to learn effective regulatory influences that are consistent with the stable transcriptional states observed in the RNA–seq data.

At the other extreme, structural priors derived from TF–gene promoter binding and protein–protein colocalization anchor inference to molecular evidence and improve interpretability. Even under the most stringent structural constraints, inferred GRNs still reproduced a substantial fraction of the observed profiles as stable states. However, enforcing priors as hard constraints reduced dynamical performance by limiting the model to regulatory interactions explicitly supported by available molecular evidence. This restriction excludes both indirect regulatory influences and direct interactions that are absent from or incompletely captured by the prior. Consequently, genes exhibiting multi–state expression and lacking evidence of direct TF binding (e.g., *SNT2*) are effectively treated as unregulated single–state genes, leading to poor fits. Generally, the incompleteness of TF binding and interaction data constrains SETIA’s ability to capture the full range of regulatory behaviors. These results support a pragmatic use of structural priors: within SETIA, priors are most effective when applied as soft constraints that guide inference without acting as absolute masks, preserving both biological interpretability and the dynamical flexibility required to reproduce and generalize transcriptional stable states.

Benchmarking GRN inference in this context highlights how different modeling strategies serve distinct goals. Under conventional edge–level evaluation, SETIA achieves higher precision and MCC than Inferelator, reflecting its prioritization of reliability over coverage. A key distinction is that SETIA infers an explicit dynamical model that can be evaluated by its ability to reproduce stable transcriptional states and generalize to unseen conditions, whereas Inferelator infers static regulatory associations and is optimized for genome–scale scalability. To bridge this gap, we implemented a semi–genome–scale strategy in which the SETIA–inferred ssTF network serves as a dynamical core, with non–TF genes incorporated as downstream targets through structurally supported edges without feedback to the core. This hybrid design extends predictions to genome–wide outputs while preserving the mechanistic and dynamical interpretability of the core regulatory module.

SETIA operates under several modeling assumptions and practical constraints that define its scope of application. First, SETIA adopts a stable–state, attractor view of gene regulation and is not designed to model transient or non–equilibrium responses, such as those observed under acute perturbations. This design simplifies the GRN model and inference process by avoiding the need to estimate detailed kinetic parameters. Second, identification of discrete gene expression states requires sufficient sampling depth. Reliable detection depends on adequate replication across conditions. Putative states supported by few samples cannot be confidently resolved computationally and require experimental validation. Third, GRN inference depends on the choice of modeled regulators, here the 76 ssTFs. This choice is practical rather than axiomatic, and there is no guarantee that regulatory control of these TFs is self–contained within this set. As a result, expression variation may reflect factors outside the modeled network or non–transcriptional mechanisms^55–57^. In unconstrained settings, such effects can be absorbed into effective edges, improving dynamical fit but reducing mechanistic specificity. Together, these considerations highlight opportunities to extend attractor–based GRN inference through improved experimental design, broader data integration, and incorporation of additional regulatory layers and dynamical regimes.

We envision SETIA’s application to multicellular organisms, where its central premise, that cell types correspond to stable states of an underlying gene regulatory system, is well suited to development and differentiation. In humans, bulk and single–cell atlases define robust transcriptional states that can serve as targets for dynamical GRN inference, enabling models whose stable states correspond to stem, progenitor, and differentiated identities. This framework naturally extends to mammalian systems, where a compact TF core can be modeled dynamically and genome–wide transcriptional outputs incorporated through structural priors such as TF–gene promoter binding and TF–TF interactions. By integrating these priors with single–cell expression data, SETIA could yield dynamical models that explain the stability of cell states and predict how perturbations reshape cell fate landscapes in development and disease.

## Supporting information

STAR Methods

## Acknowledgments

We thank Dr. Jordan Rozum for insightful advice and the members of the Pugh laboratory for helpful discussions and technical assistance. We also acknowledge the use of the following resources: the Platform for Epigenomic and Genomic Research (RRID:SCR_021861), ScriptManager (RRID:SCR_021797), Galaxy (RRID:SCR_006281), and the Pennsylvania State Institute for Computational and Data Sciences Advanced Cyberinfrastructure (RRID:SCR_025154). This work was supported by NIH R01ES034353, NIH R35GM145217 to B. Franklin Pugh. William K. M. Lai acknowledges support of NIH R35GM155380, NSF XSEDE and subsequent ACCESS award BIO220026.

## Author Contributions

R.L. conceived the study, developed the methodology, performed the experiments and data analysis, and wrote the manuscript. B.F.P. and W.K.M.L. supervised the project, contributed to study design and interpretation, and revised the manuscript.

## Supplementary Figure Legends

**Figure S1.**
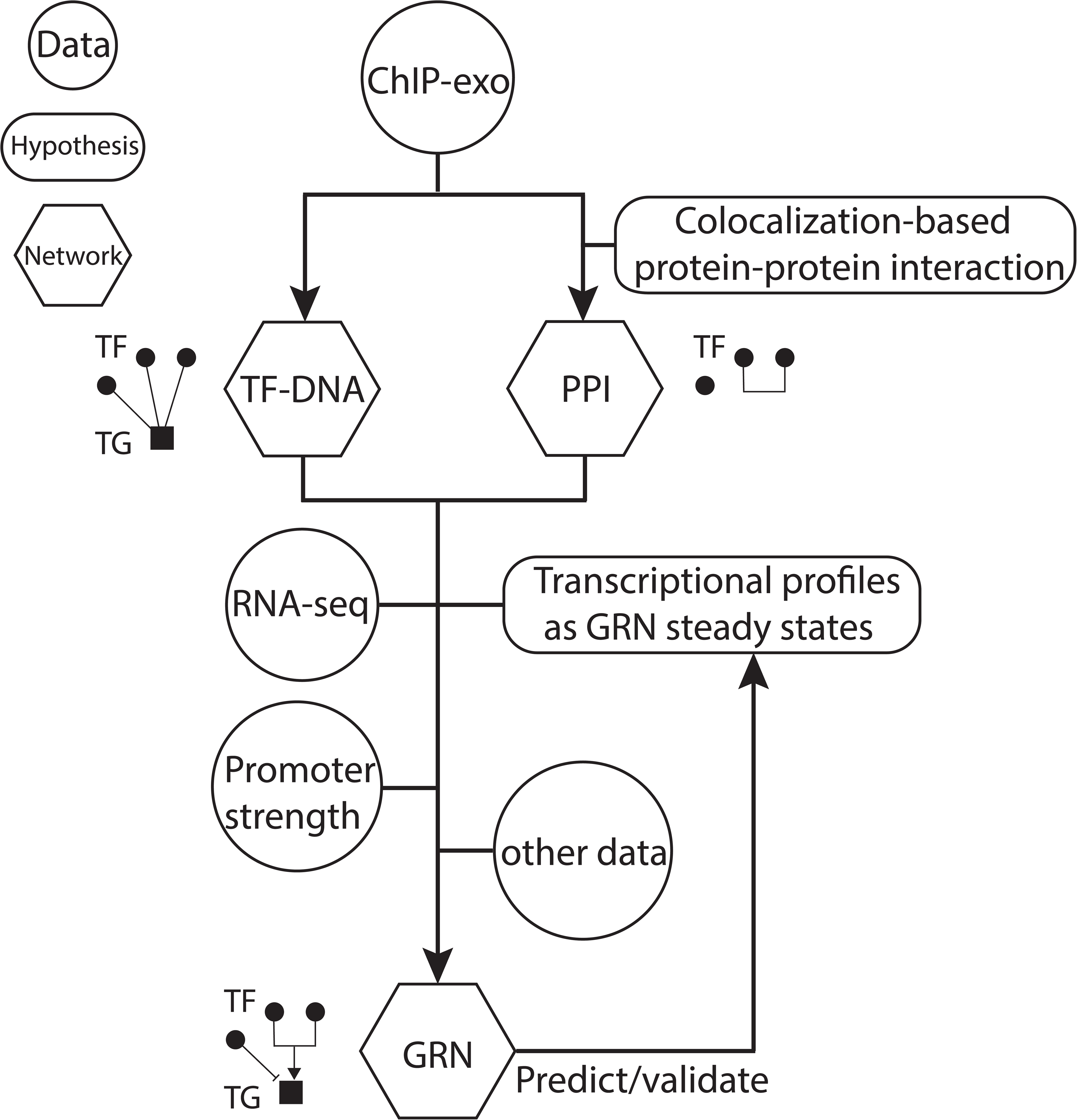
Overview of the SETIA framework for gene regulatory network inference and validation. ChIP–exo data are used to infer a TF–DNA binding network and a protein–protein interaction (PPI) network based on shared genomic binding patterns. RNA–seq profiles from perturbation experiments define discrete transcriptional stable states, while CAGE–seq provides estimation of gene–specific promoter strength. These multilayer data are integrated into an ordinary differential equation–based framework that assigns regulatory signs and optimizes kinetic parameters so that the inferred GRN reproduces observed transcriptional profiles as stable attractors, enabling prediction and validation under new conditions.

**Figure S2.**
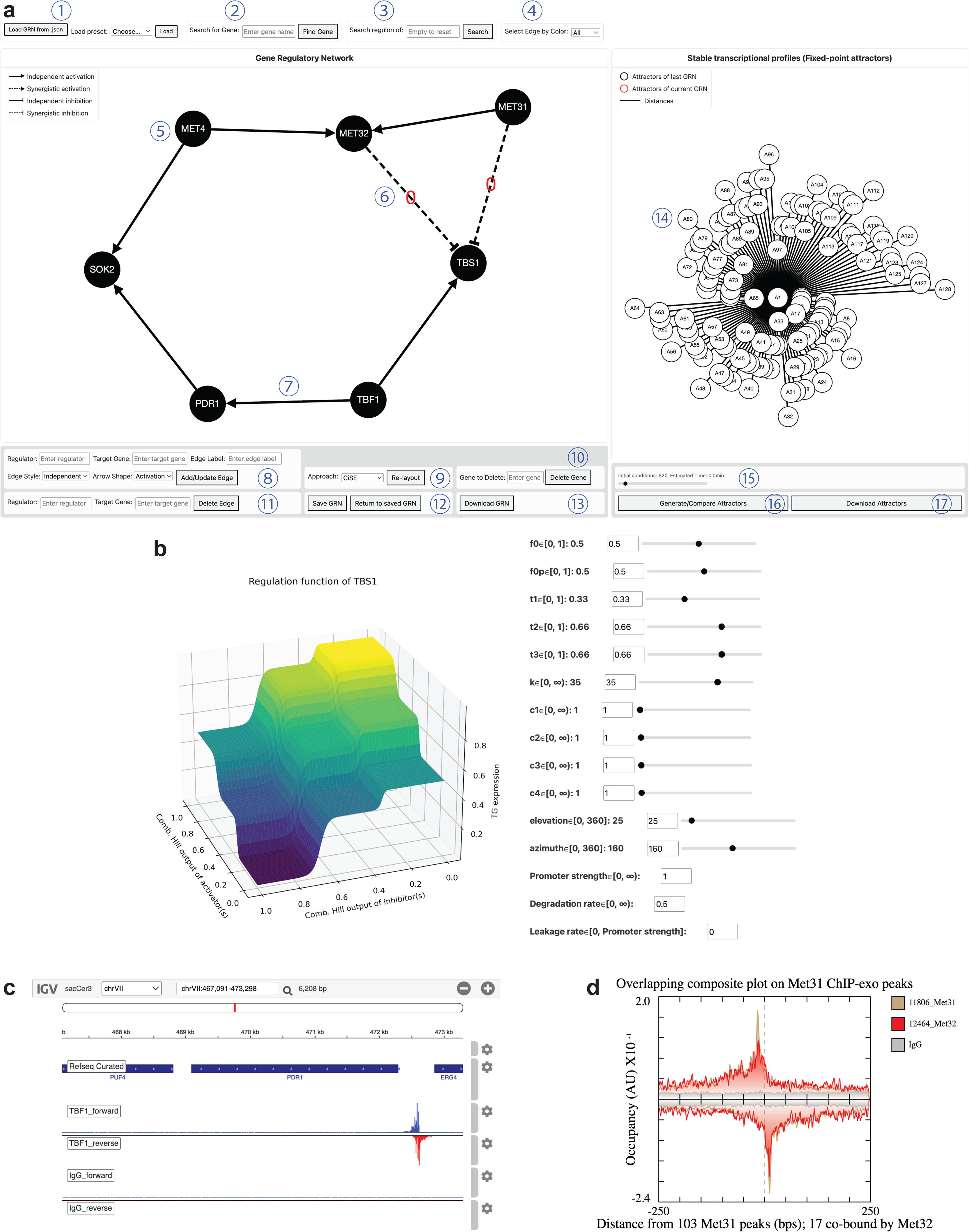
An overview of the GRN simulator website. (a) Main interface. (1) Load a network either by uploading a local JSON file or by selecting a preset network, including TF–DNA binding, protein–protein colocalization (PPC), or gene regulatory network (GRN). (2) Find a gene by name and center it in the network. (3) Subset the network to a gene of interest and its downstream targets. (4) PPC networks only: highlight edges by STRING’s confidence category (green, very high; yellow, high; orange, medium; red, exploratory). (5) Click a gene node to display its regulation function and fitted parameters. (6) Click a dashed GRN edge (synergistic regulation) to view both the IGV tracks and an overlaid ChIP–exo composite plot for the two factors, highlighting similar crosslinking patterns. (7) Click a solid GRN edge to view the ChIP–exo signal of the regulator TF at the target promoter in IGV (IgG shown as a negative–control track). (8) Add or update an edge by specifying regulator, target, an optional complex label, edge style (solid, independent; dashed, synergistic), and sign (triangle, activation; blunt, inhibition). (9) Apply a selected layout algorithm for the network. (10–12) Delete genes or edges, and save/restore a working network state. (13) Export the current network as an image and JSON. (14) Stable states (attractors) of the GRN are displayed in the right panel; edges denote Euclidean distances between attractors. (15) Generate evenly spaced initial conditions in expression space (with an estimated runtime). (16) Integrate the GRN ODE system to identify attractors. (17) Download attractor expression profiles. (b) Example regulation function and kinetic parameters for *TBS1*, editable via interactive controls. (c) Example IGV view showing ChIP–exo signal of Tbf1 at the *PDR1* promoter. (d) Example overlaid composite plots for Met31 and Met32.

**Figure S3.**
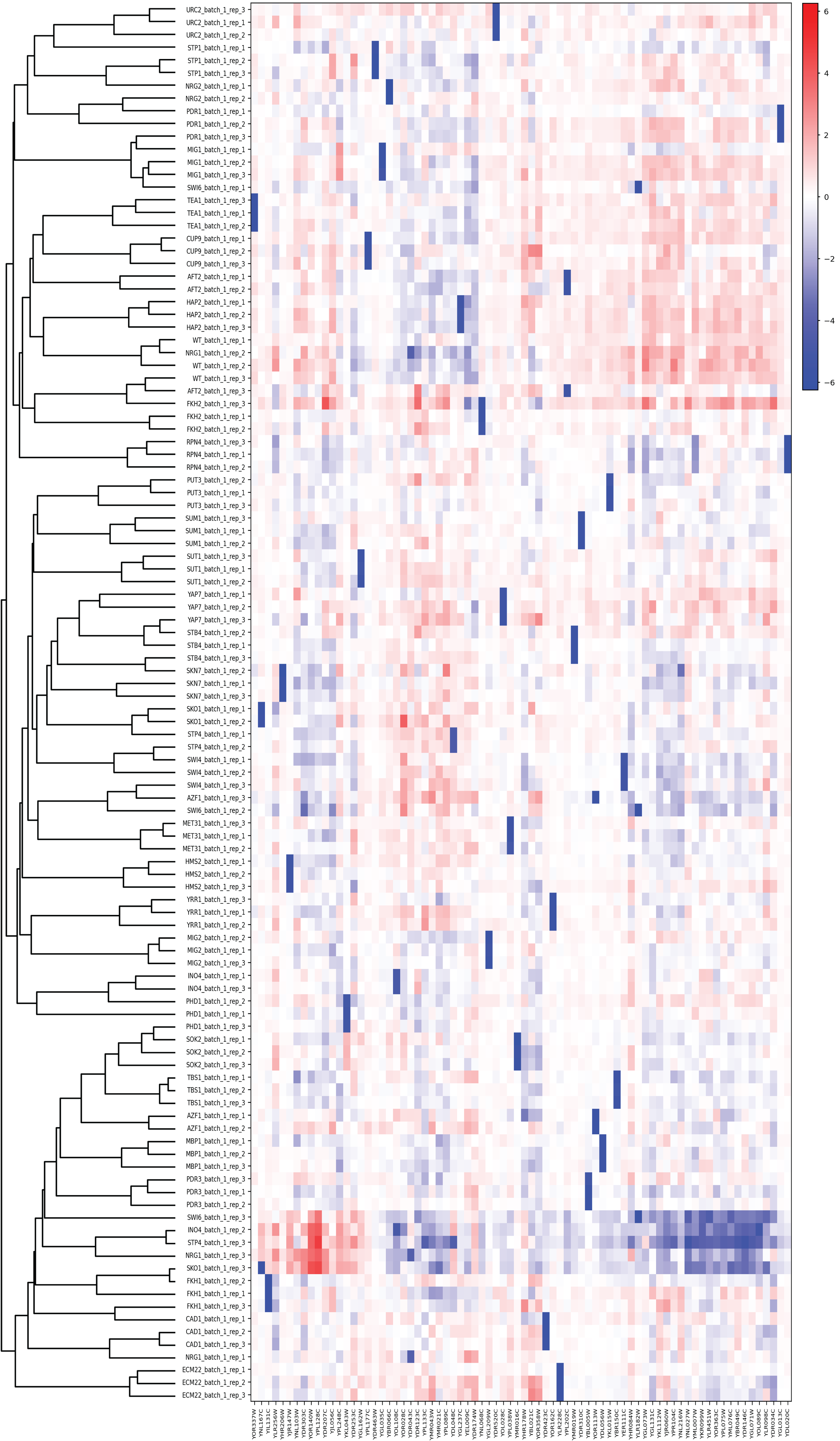
Hierarchical clustering of yeast RNA–seq samples based on the expression profiles of 76 sequence–specific transcription factor genes. Each row is one RNA–seq sample (genotype × replicate) and each column is an ssTF gene. Samples were clustered by average–linkage hierarchical clustering using correlation distance on log_2_ (CPM+1) expression values that were z–scored per gene across samples. The heatmap shows gene–wise standardized expression (blue represents lower than the gene’s mean across samples; red represents higher), and the left dendrogram indicates sample similarity.

**Figure S4.**
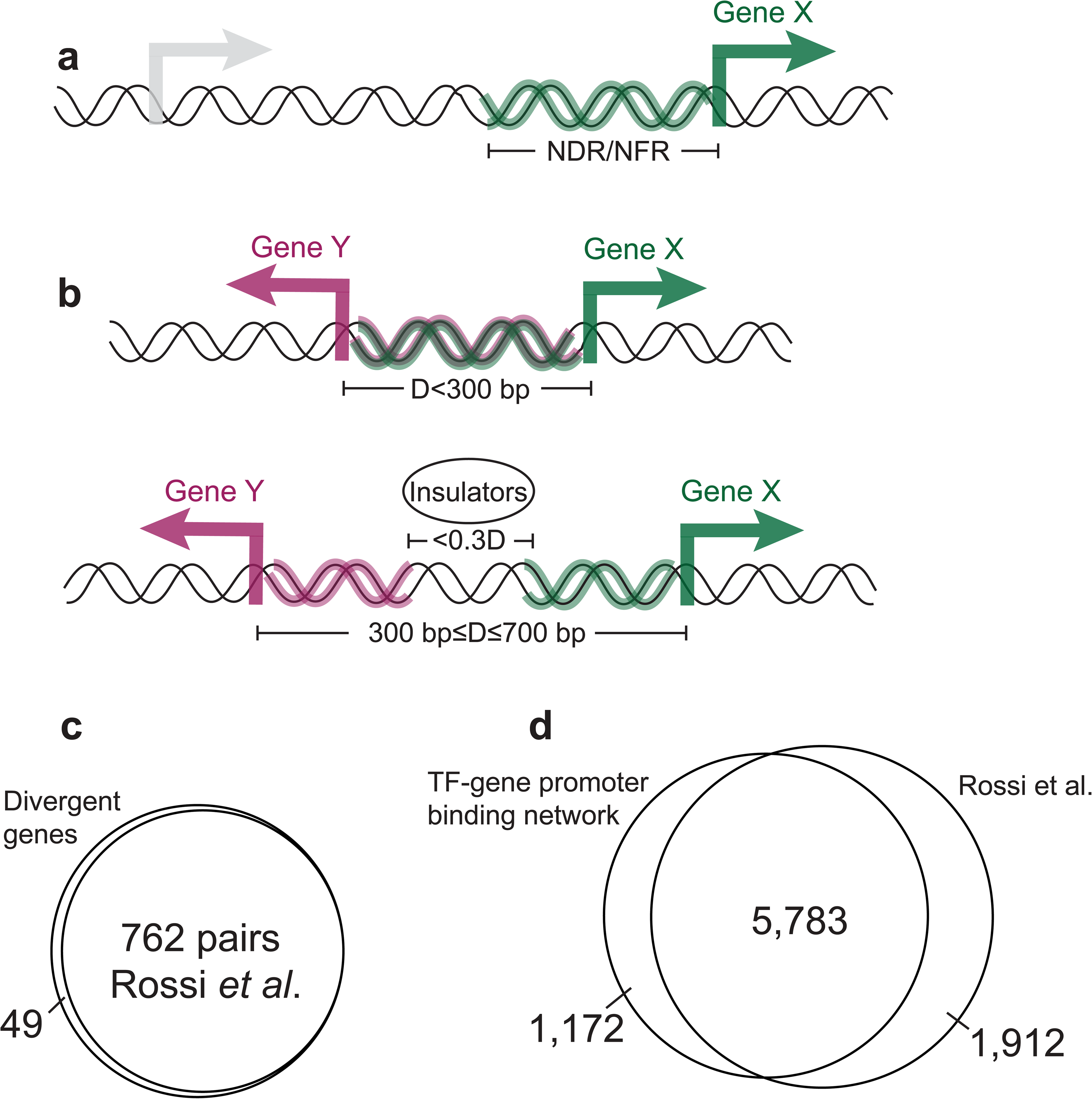
Protein–DNA binding network accounting for gene arrangement and insulators. (a) Schematic of a head–to–tail gene arrangement, showing an NDR/NFR for TF binding to the downstream target gene (Gene X). (b) Schematic of head–to–head gene pairs with transcription start sites less than 300 bp apart, allowing shared transcription factor binding to potentially influence both gene X and gene Y. For gene pairs with transcription start sites 300–700 bp apart, co–regulation may still occur unless insulator proteins (Cbf1, Rap1, or Reb1) bind in the middle within 30% of the intergenic distance, thereby blocking shared regulation. (c) Venn diagram comparing the divergent head–to–head protein–coding gene pairs identified in Rossi et al. and in this study. (d) Venn diagram comparing the protein–DNA binding networks identified in Rossi et al. and in this study.

**Figure S5.**
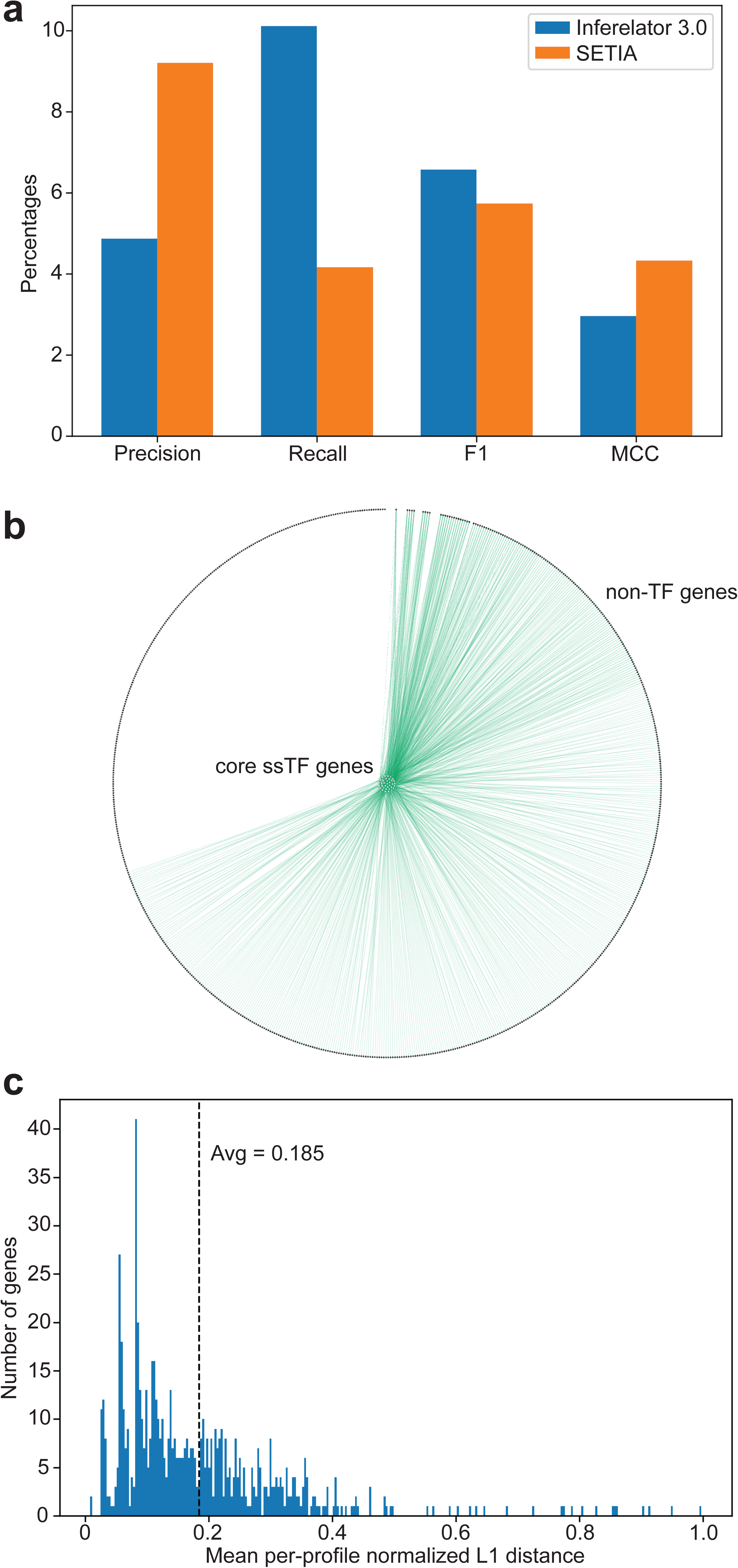
Comparison of GRN inference performance and evaluation of the semi–genome–scale model. (a) Comparison of edge–level performance metrics for GRNs inferred by Inferelator 3.0 and SETIA on the 76×76 regulator–target search space, including accuracy, precision, and recall, evaluated against a permissive reference network. (b) Semi–genome–scale GRN, with ssTF–encoding genes forming a central regulatory core and non–TF genes arranged in an outer layer. Edges from ssTFs to non–TF genes are supported by TF–DNA binding evidence and are shown in green. (c) Distribution of mean per–profile normalized L1 distances for 714 differentially expressed non–TF genes that are bound by at least one of the 76 ssTFs in the semi–genome–scale GRN. The dashed line indicates the mean error 0.185.

**Figure S6.**
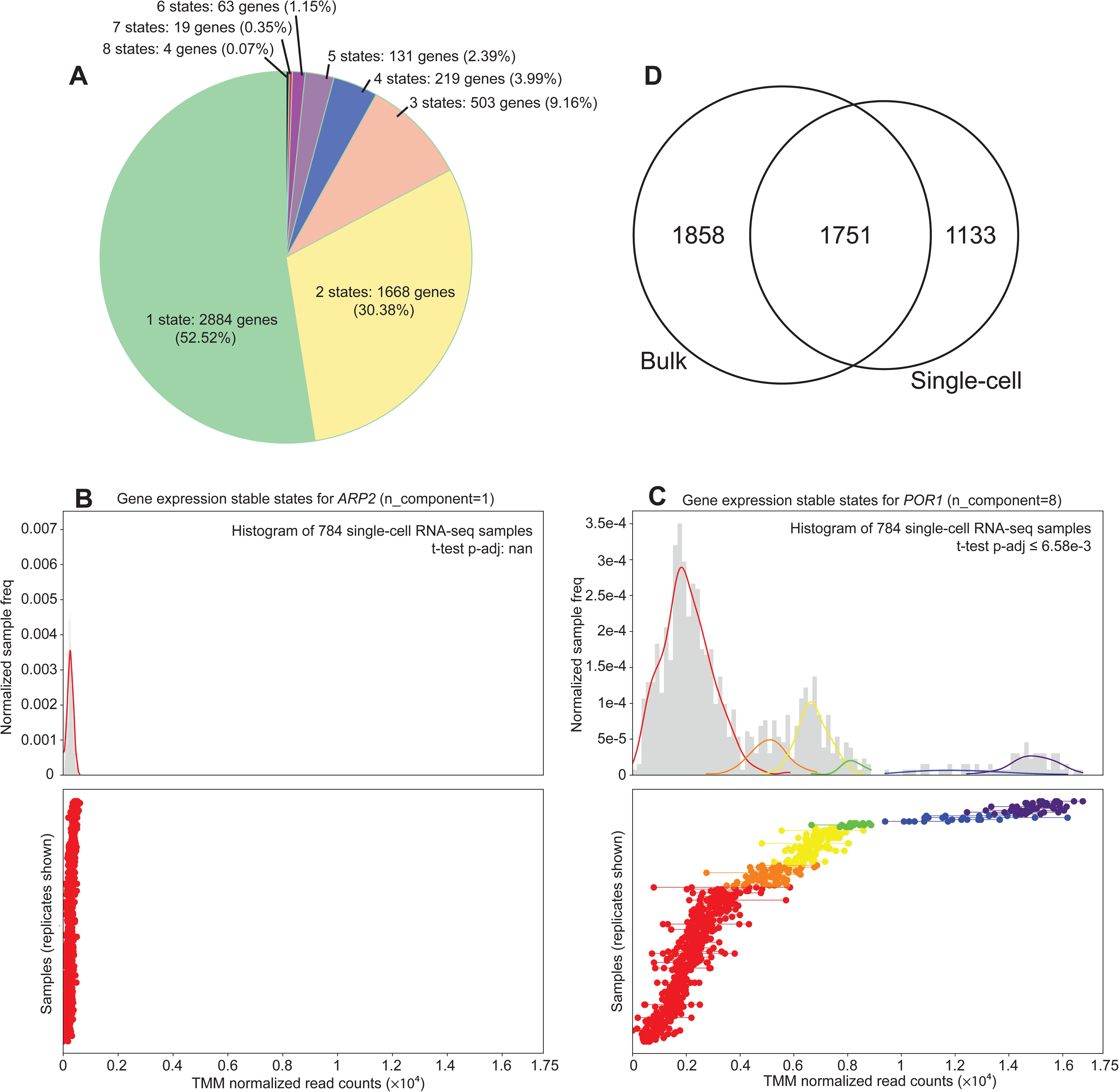
Discrete transcriptional stable states and expression variability across bulk and single–cell RNA–seq datasets. (a) Distribution of the number of discrete expression states identified per gene in the single–cell RNA–seq dataset. (b–c) Representative examples illustrating unimodal (*ARP2*) and multimodal (*POR1*) expression behavior across pseudobulk samples. Bottom panels show TMM–normalized read counts for each genotype, with rows corresponding to distinct genotypes and dots connected by lines representing biological replicates of the same genotype. Top panels summarize the same data as histograms of all samples, overlaid with kernel density estimates to highlight the number and separation of stable expression states. Underlying data and statistical details are provided in **Supplementary Data 10**. (d) Overlap of genes classified as single–state in bulk and single–cell RNA–seq datasets. Of the 3,609 genes classified as single–state in the bulk RNA–seq dataset, 1,751 (48.5%) were also classified as single–state in the single–cell dataset. Conversely, 1,751 of the 2,884 single–state genes identified in the single–cell dataset (60.7%) overlapped with those from the bulk.

**Figure S7.**
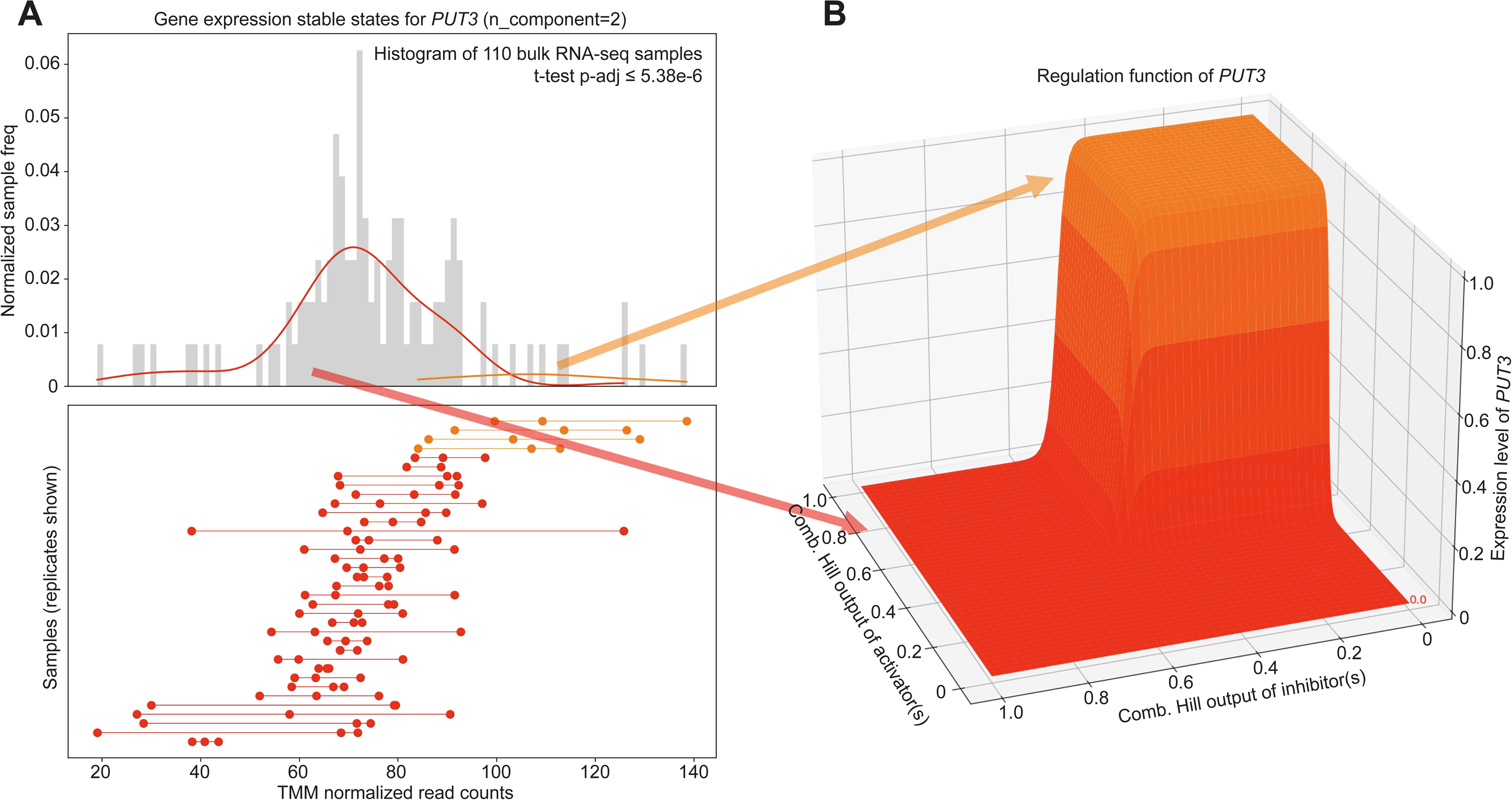
Discretization of *PUT3* expression. (a) Top: histogram of sample–level TMM–normalized Salmon read counts for *PUT3* with overlaid KDEs of the two inferred expression–state distributions. Bottom: per–genotype replicate visualization, where each dot is a replicate’s TMM–normalized count and replicates from the same genotype are connected by a line. Two stable expression states were identified; Welch’s t-test between the groups (with multiple–testing correction applied when >2 groups exist) gives an adjusted p = 5.38 x 10^-6^. (b) Regulation function derived from the discrete expression states. The orange and red plateaus indicate the means of the corresponding identified distributions and therefore the discrete regulatory levels used in downstream modeling.

**Figure S8.**
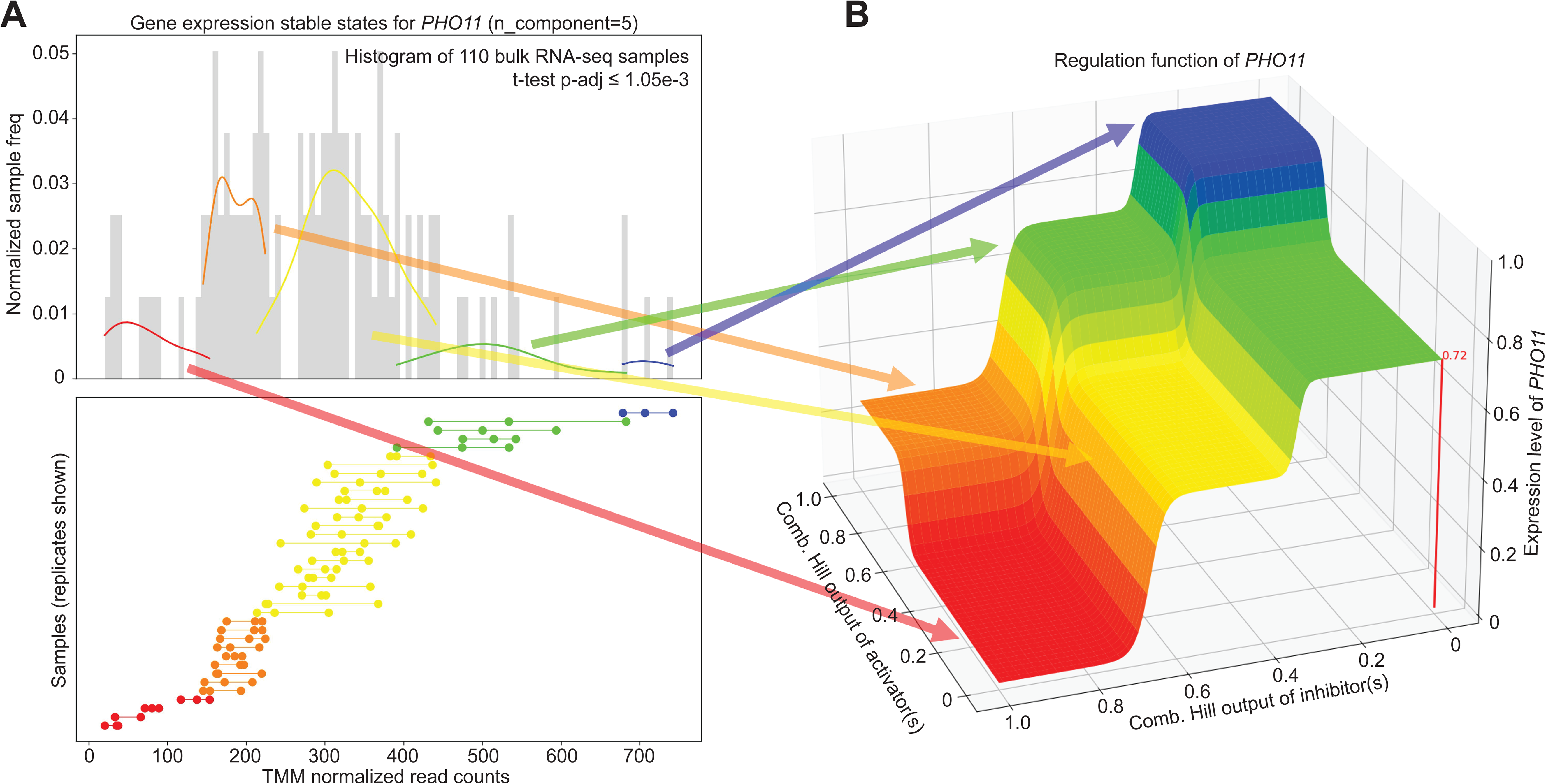
Multilevel discretization of *PHO11* expression. (a) Top: distribution of TMM–normalized Salmon read counts for *PHO11* across all samples, showing a multimodal structure captured by KDEs corresponding to five distinct expression states. Bottom: genotype–resolved replicate measurements, where each dot represents a biological replicate and lines connect replicates from the same genotype, revealing graded shifts in expression across perturbations. Five stable expression states were inferred; pairwise Welch’s t-tests between states, with multiple–testing correction, yielded a maximum adjusted p-value of 1.05 x 10^-3^. (b) Regulation function parameterized from the five discrete expression states. The terraced output illustrates multiple regulatory plateaus spanning a broad dynamic range, with each plateau corresponding to the mean expression level of one inferred state and defining a multilevel regulatory response used in downstream modeling.

## Supplementary Tables

**Supplementary Table 1.** Parameter table for the GRN dynamic system.

| Symbol | Description | Unit |
| --- | --- | --- |
| $[R]_i$ | Number of TMM normalized mRNA read counts for each gene | dimensionless |
| $V_{\max}$ | Maximal rate of transcription for each promoter | nucleotides/second |
| $V_{\min}$ | Minimal rate of transcription for each promoter | nucleotides/second |
| $f_0$ | Expression level of the target gene when no TFs act on the gene | percentage |
| $f'_0$ | Expression level of the target gene when both | percentage |
|  | activators and repressors competitively act on the gene |  |
| $t$ | Threshold constants (half-maximal concentrations) that set the positions of two distinct Hill-type response curves in the WDH function | dimensionless |
| $n$ | Hill coefficient | dimensionless |
| $c$ | A weighting factor that balances the relative contribution of the two Hill terms in the WDH function | dimensionless |
| $D_{\text{mRNA}}$ | Rate of degradation for each mRNA | 1/second |

## Supplementary Data

**Supplementary Data 1** | Inferred stable expression states and statistical details for all genes across the 37 wild–type and transcription factor knockout (TFKO) strains in the bulk RNA–seq dataset.

**Supplementary Data 2** | SETIA–inferred, structurally unconstrained gene regulatory network corresponding to **Figure 3a**.

**Supplementary Data 3** | TF–gene promoter binding network inferred from ChIP–exo data.

**Supplementary Data 4** | Protein–protein colocalization network inferred from ChIP–exo composite profiles across all TF–bound genomic locations.

**Supplementary Data 5** | Protein–protein colocalization network inferred from ChIP–exo composite profiles restricted to motif–associated TF binding sites.

**Supplementary Data 6** | SETIA–inferred gene regulatory network incorporating TF–gene promoter binding and gene–gene co–expression (absolute mutual information > 0.25).

**Supplementary Data 7** | SETIA–inferred gene regulatory network constrained exclusively by direct TF–gene promoter binding evidence.

**Supplementary Data 8** | SETIA–inferred gene regulatory network constrained by TF–gene promoter binding with additional requirement of cognate DNA motif presence at each bound site.

**Supplementary Data 9** | Full semi–genome–scale GRN inferred in this study, including regulatory interactions linking the ssTF core network to downstream non–TF genes.

**Supplementary Data 10** | Inferred stable expression states and statistical details for all genes across the wild–type and 12 transcription factor deletion strains profiled under 11 growth conditions in the single–cell RNA–seq dataset.

