## Supplementary material for "A unified framework for causal gene regulatory network inference grounded in orthogonal molecular evidence": STAR Methods

Detailed methods are provided in the online version of this paper and include the following:

### Key Resources Table

| **REAGENT or RESOURCE** | **SOURCE** | **IDENTIFIER** |
| --- | --- | --- |
| **Biological Samples** | | |
| *S. cerevisiae wild-type strain BY4741* | This study | N/A |
| *S. cerevisiae single-TF knockout strains (36 TFKO strains)* | Yeast Knockout Collection; Horizon Discovery | N/A |
| **Chemicals, Peptides, and Recombinant Proteins** | | |
| NEBNext Poly(A) Magnetic mRNA Isolation Kit | New England Biolabs | Cat# S1550S |
| NEBNext Ultra II RNA Library Prep Kit for Illumina | New England Biolabs | Cat# E7775 |
| **Deposited Data** | | |
| Bulk RNA-seq: WT and 36 TFKO strains (S. cerevisiae) | This study | GEO: GSE317148 |
| Single-cell RNA-seq: WT and 12 TF-deletion strains under 11 growth conditions | Jackson et al., 2020 | GEO: GSE125162 |
| ChIP-exo binding data (~400 genome regulatory proteins, S. cerevisiae) | Rossi et al., 2021 | Rossi et al. Nature 592, 309–314 (2021) |
| Yeast protein interactome | Michaelis et al., 2023 | Michaelis et al. Nature 624, 192–200 (2023) |
| STRING protein interaction database | Szklarczyk et al., 2023 | https://string-db.org |
| **Experimental Models: Organisms/Strains** | | |
| *Saccharomyces cerevisiae* BY4741 (MATa his3Δ1 leu2Δ0 met15Δ0 ura3Δ0) | Yeast Knockout Collection; Horizon Discovery | N/A |
| **Software and Algorithms** | | |
| SETIA (GRN inference framework) | This study | https://github.com/CEGRcode/2024_Li_GRN_inference |
| GRN simulator web interface | This study | https://grn.cac.cornell.edu:5000/ |
| Salmon v1.x (transcript quantification) | Patro et al., 2015 | https://combine-lab.github.io/salmon/ |
| ChExMix (ChIP-exo peak calling) | Yamada et al., 2019 | https://github.com/seqcode/chexmix |
| SciPy (peak detection and signal processing) | Virtanen et al., 2020 | https://scipy.org; RRID:SCR_008058 |
| ScriptManager (genomics utilities) | This study | RRID:SCR_021797 |
| Galaxy (bioinformatics platform) | Galaxy Community | RRID:SCR_006281 |
| Platform for Epigenomic and Genomic Research (PEGR) | This study | RRID:SCR_021861 |
| Inferelator 3.0 (GRN inference benchmark) | Skok Gibbs et al., 2022 | https://github.com/flatironinstitute/inferelator |
| **Other** | | |
| Pennsylvania State Institute for Computational and Data Sciences Advanced Cyberinfrastructure (ICDS-ACI) | Pennsylvania State University | RRID:SCR_025154 |

### Resource Availability

#### Lead Contact

#### Materials Availability

This study did not generate new unique reagents. Yeast knockout strains used in this study are available from the Yeast Knockout Collection (Horizon Discovery).

#### Data and Code Availability

- **RNA-seq data:** The bulk RNA-seq dataset generated in this study has been deposited in the Gene Expression Omnibus (GEO) under accession number **GSE317148** and is publicly available as of the date of publication.
- **Code:** All source code is publicly available on GitHub at https://github.com/CEGRcode/2024_Li_GRN_inference. Source code for the GRN simulator web interface is included in the same repository. DOI will be minted upon acceptance.
- **Additional information:** An online demonstration of the GRN simulator is accessible at https://grn.cac.cornell.edu:5000/. Any additional information required to reanalyze the data reported in this paper is available from the lead contact upon request.

### Experimental Model and Subject Details

#### Saccharomyces cerevisiae Strains and Culture Conditions

*S. cerevisiae* TFKO strains, together with wild-type BY4741 (*MATa his3Δ1 leu2Δ0 met15Δ0 ura3Δ0*), were obtained from the Yeast Knockout Collection (Horizon Discovery). Strains were cultured in YPD medium at 25°C and harvested at mid-log phase (OD_600_ 0.75–0.85). Cells were collected by centrifugation. A subset of 36 single-TF knockouts were used because a fraction of ssTFs is essential for viability and cannot be deleted.

### Method Details

#### RNA-seq Library Preparation and Sequencing

Total RNA was extracted from harvested yeast cells following phenol–chloroform lysis. Polyadenylated RNA was isolated using the NEBNext Poly(A) Magnetic mRNA Isolation Kit (New England Biolabs, Cat# S1550S) according to the manufacturer’s instructions, enriching for mature mRNA and reducing ribosomal RNA contamination. RNA-seq libraries were constructed from purified mRNA using the NEBNext Ultra II RNA Library Prep Kit for Illumina (New England Biolabs, Cat# E7775) and sequenced on an AVITI24 platform.

#### RNA-seq Data Processing and Transcript Quantification

Sequencing reads were aligned and transcript abundance was quantified as read counts for each ssTF-encoding gene using Salmon^58^. Read counts were normalized using the TMM (trimmed mean of M-values) method. Genes with a CPM of at least 1 in at least two samples were retained for downstream analysis. Single-cell RNA-seq expression data were obtained from Jackson et al.^59^ (GEO accession GSE125162) and processed as described in Skok Gibbs et al.^38^ The curated yeast regulatory gold standard was compiled by Tchourine et al.^39^ and processed following procedures described in Skok Gibbs et al.^38^.

**Identification of Discrete Gene Expression States**

##### Overview

To model gene expression across conditions as a mixture of discrete stable states, we implemented a hybrid, principled algorithm (Algorithm 1) to identify discrete steady-state expression modes from replicate TMM-normalized Salmon read counts. The method is built on three core principles: (1) a gene expression state is represented as a statistical distribution; (2) all biological replicates of a sample are expected to fall within the same expression state; and (3) expression states appearing in only a small subset of TFKO samples should not be disregarded even when supported by only a few samples.

##### Composite Dissimilarity Metric

A robust pairwise dissimilarity is computed between every pair of replicate groups by combining an effect-size metric (Cliff’s delta, $\delta_{ij}\in[-1,1]$) with a location-shift estimator (Hodges–Lehmann median difference, $HL_{ij}$). The composite dissimilarity is defined as:

$D_{ij}=|\delta_{ij}|\cdot|HL_{ij}|.$ (Equation 1)

This composite measure captures both consistency and magnitude of separation between distributions and is robust to non–Gaussian shapes and unequal sample sizes.

##### Elbow-Driven Nonparametric Merging

Composite dissimilarity values are swept over a fine grid (default 100 steps) to identify connected components via union-find at each threshold. The elbow in the number-of-clusters-versus-cutoff curve determines an optimal merging threshold$\tau^{*}$. Single-sample clusters are filtered after initial merging.

##### Greedy Parametric Consolidation

Resulting clusters are further consolidated using a greedy Welch’s t-test merger. At each step, pairwise Welch p-values are computed between all cluster pairs (requiring ≥2 observations per cluster). An empirical KDE-based permutation likelihood augments the p-value where applicable ($p_{ab}\leftarrow max(p_{ab},\text{KDE\_likelihood})$). The cluster pair with the largest (least significant) p-value is merged if $p\geq\alpha$ (default $\alpha=0.01$). This two-stage approach reduces sensitivity to outliers while allowing statistically defensible merges.

**Algorithm 1**: Identification of discrete gene expression states

**Require:** Expression vectors for one gene from all biological replicates $\{\mathbf{x}_{1},\ldots,\mathbf{x}_{R}\}$ (TMM–normalized counts);
1:	grid size $M$ (default $100$); Welch merge threshold $\alpha$ (default ${10}^{-2}$).

**Ensure:** Final clusters (gene expression stable states) $\{\mathcal{C}_{1},\ldots,\mathcal{C}_{K}\}$; sample–to–state mapping; per–state summaries (median, mean, std).

2:	**(A) Preliminaries**

3:	Compute per-replicate within–replicate variability:

$$v_{r}=\left\{ \begin{matrix} 0 & \text{if }|\mathbf{x}_{r}|<2, \\ \frac{1}{\binom{n_{r}}{2}}\sum_{i<j} |x_{r,i}-x_{r,j}| & \text{otherwise.} \end{matrix} \right.$$

4:	Flatten values for later use: $V=\bigcup_{r}\mathbf{x}_{r}$.

5:	**(B) Pairwise composite dissimilarity**

6:	**for all** replicate pairs $(i,j)$ **do**

7:	compute Cliff’s delta $\delta_{ij}\in[-1,1]$ between $\mathbf{x}_{i},\mathbf{x}_{j}$

8:	compute Hodges–Lehmann shift $HL_{ij}=median(\{x-y:x\in\mathbf{x}_{i},y\in\mathbf{x}_{j}\})$

9:	define composite dissimilarity

$$D_{ij}=|\delta_{ij}|\cdot|HL_{ij}|.$$

10:	**end for**

11:	Build pairwise table $\mathcal{T=\{(}i,j,D_{ij})\}$.

12:	**(C) Elbow–driven nonparametric merging**

13:	Let $\{\tau_{1},\ldots,\tau_{M}\}$ be $M$ values evenly spanning the observed $D_{ij}$ range.

14:	**for** $m\leftarrow1 \mathrm{to}M$ **do**

15:	form graph $G_{m}$ on replicate indices where edge $(i,j)$ present if $D_{ij}\leq\tau_{m}$

16:	compute connected components of $G_{m}$ via union–find $\Rightarrow$ $K(\tau_{m})$ clusters

17:	**end for**

18:	Determine elbow index $m^{*}$ of $K(\tau)$ (largest, left–most drop that falls below a cutoff criterion); set $\tau^{*}=\tau_{m^{*}}$.

19:	Form initial clusters $\{\mathcal{C}_{k}\}$ by merging groups connected at threshold $\tau^{*}$.

20:	Remove clusters with fewer than two observations (singletons treated separately).

21:	**(D) Greedy parametric consolidation (Welch)**

22:	Initialize merged clusters $\tilde{\mathcal{C}}\leftarrow\{\mathcal{C}_{k}\}$ and mapping to original replicate indices.

23:	**while** True **do**:

24:	Compute pairwise Welch $t$-test $p$-values $p_{ab}$ between all cluster pairs $({\tilde{\mathcal{C}}}_{a},{\tilde{\mathcal{C}}}_{b})$ (require $\geq2$ obs per cluster).

25:	Where available, augment $p_{ab}$ with an empirical KDE permutation likelihood to increase robustness; set $p_{ab}\leftarrow max(p_{ab},\text{KDE\_likelihood})$.

26:	Let $(i,j)=arg\max_{a\neq b}p_{ab}$ and $p_{\max}=p_{ij}$.

27:	**if** $p_{\max}<⍺$ **or** no valid pairs remain **then**

28:	**break**

29:	**else**

30:	Merge cluster $j$ into $i$ (concatenate observations and merge original–index mappings); remove cluster $j$.

31:	**end if**

32:	**end while**

33:	**(E) Finalization and summaries**

34:	**for** each final cluster ${\tilde{\mathcal{C}}}_{k}$ **do**

35:	compute robust summaries: median $\tilde{\mu}_{k}$, mean $\bar{\mu}_{k}$, and within–cluster standard deviation $\sigma_{k}$

36:	compute weight $w_{k}=|{\tilde{\mathcal{C}}}_{k}|/|V|$

37:	**end for**

38:	Assign each biological sample to a state by mapping its replicate index to the final cluster (use the stored mapping).

39:	**return** final clusters $\{{\tilde{\mathcal{C}}}_{k}\}$, sample assignments, and summaries $\{\tilde{\mu}_{k},\bar{\mu}_{k},\sigma_{k},w_{k}\}$.

#### In Silico Benchmarking of Discrete Expression State Identification

To benchmark Algorithm 1 under controlled conditions with known ground truth, synthetic gene expression datasets were generated as finite mixtures of Gaussian expression states, with all biological replicates of a given condition constrained to arise from the same latent state. State separability was parameterized by requiring that the minimum pairwise distance between state means satisfy $|\mu_{i}-\mu_{j}|\geq s\cdot\sqrt{\sigma_{i}^{2}+\sigma_{j}^{2}}$, where $s$ is a tunable separability parameter. Intuitively, for two equal–variance Gaussian distributions, $s=0$ corresponds to complete overlap (100%), whereas $s=1,2,3$ reduce overlap to approximately 48%, 15.7%, and 3.4%, respectively. Replicate groups were assigned to states uniformly at random and expression values were sampled from the corresponding normal distribution. To ensure non–negative expression values, samples with negative values were rejected and resampled.

##### Accuracy of Discrete Gene Expression State Identification on Synthetic Data

Algorithm 1 was applied to synthetic datasets across a range of adjusted p-value cutoffs to infer discrete expression states. For each gene, predicted state labels were compared to ground-truth labels assigned during simulation. Accuracy for each gene was defined as the fraction of replicate groups whose inferred state matched the ground-truth state after label alignment. Overall accuracy at a given cutoff was computed by averaging across all simulated genes. Performance was evaluated across multiple separability regimes to assess robustness as states became increasingly overlapping (**Figure 2e**).

#### Building the Protein–Protein Colocalization (PPC) Network

##### ChIP-exo Composite Profile Analysis

ChIP–exo binding patterns of ~400 genome regulatory proteins were analyzed by centering on peaks identified by ChExMix^41^ and stratifying sites based on whether the ChExMix peak contained the cognate DNA motif of the assayed factor. Two complementary metrics were used to quantify similarity between ChIP–exo composite profiles.

##### Jensen–Shannon Divergence Distance (JSDD)

The JSDD measures global dissimilarity between normalized crosslinking distributions of two factors and is sensitive to differences in peak position and shape (**Figure 4a**). It is defined as:

$JSDD(P\parallel Q)=\sqrt{\frac{1}{2}\sum_{i} P(i)log\frac{2P(i)}{P(i)+Q(i)}+\frac{1}{2}\sum_{i} Q(i)log\frac{2Q(i)}{P(i)+Q(i)}},$ (Equation 2)

A JSDD of 0 indicates identical profiles; a value of 1 indicates completely dissimilar profiles. Potential cofactors were identified by locating the first extremum in the second derivative of the JSDD distribution, corresponding to a natural separation between closely matching and dissimilar profiles (**Figure 4b**).

##### K Ratio

The K ratio quantifies relative signal scaling at corresponding peak regions independent of positional differences (**Figure 4a**). Peaks were first identified in the queried protein’s composite profile using SciPy^60^, and their locations and prominences determined. The K ratio is defined as the scalar that minimizes the prominence-weighted Euclidean distance between the average ChIP–exo signal within peaks of the queried protein and the corresponding regions in a candidate colocalizing factor.

##### Secondary Confidence Sorting

To further refine candidate cofactors, a secondary sorting score was computed as the product of peak prominence, width, and complexity of the composite profile (**Figure 4c**). Profile complexity was quantified as the area under spikes in a y-coordinate–value frequency plot: a larger area indicates more repeating y-values and therefore a less complex (lower-confidence) profile. Factors meeting both JSDD and K ratio criteria and passing secondary sorting were designated as high-confidence colocalizing proteins.

##### Network Construction

Two PPC networks were constructed: one based on all TF-bound genomic locations, and one restricted to motif-associated binding sites (**Supplementary Data 4** and **Supplementary Data 5**). Overlap with the yeast protein interactome^48^ and the STRING database^49^ was assessed at four confidence levels (very high, high, medium, and exploratory). The union of both PPC networks and STRING interactions classified as “very high confidence” was used as the TF–TF interaction prior for GRN inference.

#### Construction of the TF–Gene Promoter Binding Network

The TF–gene promoter binding network was constructed by integrating site-specific TF–DNA binding data from ChIP–exo experiments^34^ with promoter architecture annotations, including nucleosome-depleted or nucleosome-free regions (NDRs/NFRs), promoter orientation (divergent or tandem), and the presence or absence of insulator proteins (Abf1, Rap1, or Reb1). For tandem (head-to-tail) gene configurations, a TF–target interaction was assigned when a ChIP–exo peak was detected within the target gene’s promoter NDR/NFR (**Figure S4a**). For divergent (head-to-head) gene pairs, co-regulation was assumed when the intergenic distance was <300 bp, or when the distance was 300–700 bp and no insulator protein was bound within 30% of the intergenic region (**Figure S4b**). Pairs with intergenic distances >700 bp were not considered co-regulated. This analysis identified 811 divergent gene pairs with shared bidirectional promoters, including all 762 pairs reported by Rossi et al.^34^. The resulting network contains 5,864 nodes and 6,955 edges (**Supplementary Data 3**).

#### GRN Model Architecture

##### Regulatory Edge Types

GRNs consist of nodes representing genes and the TFs they encode, and directed edges representing regulatory interactions. Edges are classified as *solid* (independent regulation, where a single TF directly regulates a target gene without involvement of other factors) or *dashed* (cooperative regulation, where multiple TFs act jointly on a target). Multiple dashed edges sharing the same numeric label converge synergistically on a target gene. For example, Met31 and Met32 both have dashed edges converging on *TBS1*, indicating cooperative inhibition of its expression (**Figure S2a**). A list of symbols and parameters used in the GRN dynamic system is provided in **Supplementary Table 1**.

##### Regulation Function

Each target gene’s expression is modeled using a combinatorial Hill function. For $m$ synergistic regulator groups, define$G_{j}(\mathbf{x})=\prod_{i\in S_{j}} x_{i}$, where $S_{j}$is the set of regulators acting jointly in group $j$. The normalized combinatorial Hill function is:

$H_{\mathrm{comb}}(\mathbf{x};\{\tau_{j}\},n) = \frac{(\sum_{j=1}^{m} \frac{G_{j}(\mathbf{x})}{\tau_{j}})^{n}}{(\sum_{j=1}^{m} \frac{G_{j}(\mathbf{x})}{\tau_{j}})^{n}+1},$	(Equation 3)

where $x_{i}$ denotes the TMM–normalized Salmon read count of regulator $i$, $\tau_{j}$ is the half–saturation threshold for group $j$, and $n$ is the Hill coefficient that controls steepness. To capture three discrete effects on a target gene, a Weighted Dual-Hill (WDH) function is constructed by combining two combinatorial Hill functions with thresholds $t_{1}$ and $t_{2}$ and a weight factor $c$:

$WDH(\mathbf{x};t_{1},t_{2},n,c)=\frac{1}{1+c}\left( H_{\mathrm{comb}}(\mathbf{x};t_{1},n)+c H_{\mathrm{comb}}(\mathbf{x};t_{2},n) \right).$ (Equation 4)

Activators and repressors enter the model through separate WDH terms. Let $[R]_{act}$ and $[R]_{rep}$ denote activator and repressor expression vectors, respectively. The net regulation function is:

$\begin{aligned} f_{\mathrm{GRN}}([R];\Theta)&=f_{0}+(1-f_{0}) WDH([R]_{act};t_{1},t_{2},n,c_{1})-f_{0} WDH([R]_{rep};t_{1},t_{3},n,c_{3}) \\ & +\left( f_{0}^{'}+f_{0}-1 \right) WDH([R]_{act};t_{1},t_{2},n,c_{2}) WDH([R]_{rep};t_{1},t_{3},n,c_{4}), \end{aligned}$	(Equation 5)

where $[R]$ collects the regulator variables and $\Theta$ denotes all remaining parameters. The parameter $f_{0}$ sets basal expression when regulator input is absent and $f_{0}^{'}$ sets the expression when both activators and repressors are maximal. Equation 5 enforces intuitive boundary conditions: each WDH term is normalized to [0,1]; the activator term appears with a positive weight and the repressor term with a negative weight, ensuring monotone responses. The multiplicative cross-term allows non-additive interactions between activators and repressors, permitting a distinct joint plateau when both are present.

To connect the regulation function to discrete stable expression states, plateau heights are fit in two steps. First, stable expression levels are inferred from empirical read counts using Algorithm 1, with component means providing target plateau heights. Second, parameters $f_{0}$, $f_{0}^{'}$, thresholds $t$, and weighting factors $c$ are fit via least-squares regression. The Hill coefficient n is estimated by matching the empirical kernel density estimate of observed expression to the distribution predicted by the regulation function across observed regulator values. The optimal n is often large and is capped at 30. Plateau heights are scaled so the lowest and highest plateaus correspond to 0 and 1 (**Figure S7b**; **Figure S8b**).

##### GRN Kinetic System

The GRN is formalized as a system of ordinary differential equations (ODEs). Promoter strength ($V_{\max}$, maximal transcription rate; $V_{\min}$, leak rate) was estimated from CAGE-seq or nascent RNA-seq data^27^. The ODE for gene $i$ is:

$\frac{d [R]_{i}}{dt}=V_{\min}+\left( V_{\max}-V_{\min} \right) f_{\mathrm{GRN}}\left( [R]_{*};\Theta\right)-D_{\mathrm{mRNA}} [R]_{i},$ (Equation 6)

where $[R]_{*}$ are the regulator expression levels and $D_{\mathrm{mRNA}}$ is the first-order mRNA degradation rate. A state is called stable when: (1) $|d[R]/dt|$ falls below a threshold (the larger of 1% of the gene’s maximum expression and an absolute value of 0.5); and (2) the linear stability condition holds: $\frac{d}{d[R]}(d[R]/dt)$ < 0 at that point. These two requirements together ensure that identified states are true stable fixed points.

#### GRN Inference by SETIA

##### Integrating Multiple Independent Omics Datasets

To infer a biologically coherent GRN, SETIA integrates complementary omics datasets that together constrain both network structure and kinetics. TF–DNA specificity is required by ChIP–exo peak enrichment within nucleosome-free promoter regions. Protein–protein colocalization patterns in genome-wide ChIP–exo composite binding profiles support TF–TF interactions. The stable states produced by the GRN kinetic model must recapitulate transcriptional profiles observed across diverse conditions including knockout strains, with promoter strengths estimated from CAGE-seq or nascent RNA-seq imposing dynamical constraints on synthesis rates. Satisfying these multi-layered criteria enforces mechanistic consistency across binding, interaction, and regulatory dimensions.

##### Sub-Network Decomposition and Parallel Inference

Starting from TF–DNA binding and TF–TF interaction priors, SETIA constructs the initial GRN structure (**Figure 1a**), in which each TF–target relationship can be assigned as activation, repression, or no interaction. TF–DNA binding priors constrain the search space by permitting activation or repression only for TFs with promoter-binding evidence at the target gene, while unsupported TF–target pairs are fixed as no interaction. TF–TF interaction priors restrict which TFs may be modeled as cooperative regulators of the same target gene. The constrained GRN is then decomposed into independent target-specific sub-networks, each governing expression dynamics of a single gene (**Figure 1b**). Decomposition reduces computational complexity and permits parallel numerical integration of the ODEs for each component.

##### Model Fit, Convergence, and Iterative Refinement

Transcriptomic profiles serve as initial states for each sub-network, which is simulated forward in time to identify corresponding stable states. Model fit is quantified by an element-wise, range-normalized L1 distance between initial and final states:

$D = \frac{1}{M N}\sum_{j=1}^{M} \sum_{i=1}^{N} \left| \frac{[R]_{i,j}^{\mathrm{init}}-[R]_{i,j}^{\mathrm{final}}}{\max_{1\leq k\leq M} [R]_{i,k}^{\mathrm{init}}-\min_{1\leq k\leq M} [R]_{i,k}^{\mathrm{init}}} \right|,$	(Equation 7)

where $N$ is the number of genes and M is the number of samples. This metric scales each gene by its observed dynamic range before averaging over genes and samples.

During iterative refinement, all previously tested sub-network configurations are recorded to avoid redundant simulations. A sub-network is retained once kinetic training converges, defined as the L1 distance falling below a user-defined threshold (**Figure 1c**). Sub-networks that do not meet the convergence criterion continue to be revised until they succeed or the maximum number of refinement iterations is reached. To constrain the combinatorial search space, refinement steps allow existing putative protein–protein complexes to be decomposed into smaller components, but prohibit the formation of new or larger complexes not supported by the initial prior. When all components have converged or the iteration limit is reached, converged sub-networks are reassembled into the final integrated GRN (**Figure 1d**).

##### Structural Prior Variants

Four GRN variants were inferred with progressively stricter structural constraints: (A) no structural priors; (B) TF–gene promoter binding OR co-expression (absolute mutual information > 0.25) (**Supplementary Data 6**); (C) TF–gene promoter binding only from ChIP–exo evidence (**Supplementary Data 7**); (D) TF–gene promoter binding with additional requirement of cognate DNA motif presence at the bound site (**Supplementary Data 8**). For cross-validation, the 37 transcriptional profiles were randomly partitioned into five non-overlapping subsets; GRNs were inferred from the remaining profiles and evaluated on held-out profiles.

#### Semi-Genome-Scale GRN Extension

To extend SETIA beyond the 76 ssTFs, the ssTF GRN was treated as a dynamical core and non-TF genes were connected as downstream targets through direct regulatory edges supported by TF–DNA binding evidence. Non-TF genes do not feed back into the core network. This semi-genome-scale GRN was evaluated on 714 genes that exhibited more than one discrete expression state in the bulk RNA-seq data and were bound by at least one of the 76 ssTFs (**Figure S5b**; **Figure S5c**). The full semi-genome-scale GRN is provided as **Supplementary Data 9**.

#### GRN Benchmarking Against Inferelator 3.0

Inferelator 3.0^38^ was applied to the same pseudobulk single-cell dataset in a prior-free setting; regulatory edges were selected using the automatically determined confidence cutoff (≥0.3788). The inference was restricted to 76 × 76 = 5,776 possible directed regulator–target edges among the ssTFs. Because filtering existing gold standard networks^38,39^ to include only interactions among these 76 TFs yields only 14 annotated edges, a permissive reference set was defined as the union of the curated gold standard^39^ and the TF–gene promoter binding network constructed in this study (168 positive edges; 5,608 negative edges). Performance was assessed using precision, recall, F1 score, and Matthews correlation coefficient (MCC) (**Figure S5a**).

### Quantification and Statistical Analysis

Gene expression states were identified using the two-stage procedure described above (Algorithm 1; default $\alpha=0.01$ with Bonferroni correction for multiple pairwise comparisons). All pairwise state comparisons used Welch’s t-test with multiple-testing correction; the largest adjusted p-value across all state pairs is reported in the relevant figures. GRN dynamical performance was quantified using the mean per-gene range-normalized L1 distance (Equation 7) between initial and final expression vectors across all transcriptional profiles. Statistical comparisons between GRN variants and random controls (fully random and sparsity-matched) were assessed by comparing distributions of L1 distances. Benchmarking metrics (precision, recall, F1, MCC) were computed on a 5,776-edge search space with 168 positives (<3%). The number of samples (n) evaluated for each GRN is indicated in the relevant figures.

### Additional Resources

An interactive GRN simulator for visualization, perturbation, and dynamical analysis of inferred networks is available at https://grn.cac.cornell.edu:5000/. The simulator supports loading of TF–DNA binding networks, PPC networks, and GRNs in JSON format; interactive exploration of regulation functions and kinetic parameters; IGV visualization of ChIP–exo signals at target promoters; and simulation of GRN attractors from arbitrary initial states. An overview of the simulator interface and usage guidelines is provided in **Figure S2**.
